# Cannabinoid receptor 1 positive allosteric modulator ZCZ011 shows differential effects on behavior and the endocannabinoid system in HIV-1 Tat transgenic female and male mice

**DOI:** 10.1101/2024.05.10.593514

**Authors:** Barkha J Yadav-Samudrala, Hailey Dodson, Shreya Ramineni, Elizabeth Kim, Justin L Poklis, Dai Lu, Bogna M Ignatowska-Jankowska, Aron H Lichtman, Sylvia Fitting

**Affiliations:** Department of Psychology and Neuroscience, University of North Carolina at Chapel Hill, Chapel Hill, North Carolina, USA; Department of Pharmacology and Toxicology, Virginia Commonwealth University, Richmond, Virginia, USA; Department of Pharmaceutical Sciences, Texas A&M, College Station, Texas, USA; Neoronal Rhythms in Movement Unit, Okinawa Institute of Science and Technology, Okinawa, Japan

## Abstract

The cannabinoid receptor type 1 (CB_1_R) is a promising therapeutic target for various neurodegenerative diseases, including HIV-1-associated neurocognitive disorder (HAND). However, the therapeutic potential of CB_1_R by direct activation is limited due to its psychoactive side effects. Therefore, research has focused on indirectly activating the CB_1_R by utilizing positive allosteric modulators (PAMs). Studies have shown that CB_1_R PAMs (ZCZ011 and GAT211) are effective in mouse models of Huntington’s disease and neuropathic pain, and hence, we assess the therapeutic potential of ZCZ011 in a well-established mouse model of neuroHIV. The current study investigates the effect of chronic ZCZ011 treatment (14 days) on various behavioral paradigms and the endocannabinoid system in HIV-1 Tat transgenic female and male mice. Chronic ZCZ011 treatment (10 mg/kg) did not alter body mass, locomotor activity, or anxiety-like behavior regardless of sex or genotype. However, differential effects were noted in hot plate latency, motor coordination, and recognition memory in female mice only, with ZCZ011 treatment increasing hot plate latency and improving motor coordination and recognition memory. Only minor effects or no alterations were seen in the endocannabinoid system and related lipids except in the cerebellum, where the effect of ZCZ011 was more pronounced in female mice. Moreover, AEA and PEA levels in the cerebellum were positively correlated with improved motor coordination in female mice. In summary, these findings indicate that chronic ZCZ011 treatment has differential effects on antinociception, motor coordination, and memory, based on sex and HIV-1 Tat expression, making CB_1_R PAMs potential treatment options for HAND without the psychoactive side effects.

## Introduction

According to the World Health Organization, in 2022, there were approximately 39 million people infected with human immunodeficiency virus type-1 (HIV-1) globally [1]. The development of combined antiretroviral therapies (cART) over the past four decades has significantly increased the life expectancy of people living with HIV (PLWH) [2], and therefore, it is no longer considered a death sentence but a chronic disease [3, 4]. However, as cART does not eradicate the virus, and due to low brain penetration of cART across the blood-brain barrier, low levels of viral replication and chronic immune activation still linger [5]. As a consequence, PLWH on cART display synaptodendritic damage underlying the neurocognitive impairments known as HIV-associated neurocognitive deficits (HAND) [6, 7]. Although the virus does not directly infect neurons, neural damage and injury occurs indirectly through the toxic substances from infected microglia, viral proteins, cytokines/chemokines, and free radicals [8, 9, 10]. The HIV-1 transactivator of transcription (Tat) protein is a neurotoxin that plays a major role in the pathogenesis of HAND [11, 12, 13, 14, 15]. It is known that Tat directly dysregulates the α-amino-3-hydroxy-5-methyl-4-isoxazolepropionic acid / *N*-methyl-*d*-aspartate (AMPA/NMDA) receptor causing increased intracellular sodium and calcium leading to enhanced cellular excitation and loss of dendritic structures [16, 17, 18, 19, 20, 21]. Additionally, Tat promotes neuroinflammatory signaling [22, 23, 24, 25], which is a critical component of HAND pathogenesis [26]. The persistence of HAND in the cART era provokes questions about the potential treatment of HIV-1-related brain disorders and whether cognitive deficits are reversible [7]. One avenue for treating HAND, that has recently received a lot of attention, involves the modulation of the endocannabinoid system.

The cannabinoid receptor type-1 (CB_1_R) is the most abundant G-protein coupled receptor in the central nervous system (CNS) that is mainly expressed in neurons and astrocytes [27, 28]. CB_1_Rs on neurons primarily modulate neurotransmission, whereas CB_1_Rs on astrocytes regulate intracellular calcium and facilitate neural communication [29, 30]. Endogenous cannabinoids [*N*-arachidonoylethanolamine (anandamide, AEA) and 2-arachidonoylglycerol (2-AG)] are released on demand in response to various stimuli and exert their effects mainly via CB_1_Rs [31]. Numerous studies have shown that CB_1_R activation offers neuroprotective effects by inhibiting the synaptic release of glutamate [32, 33, 34, 35] and downregulating NMDA receptor activity [36, 37, 38, 39, 40]. Further, endocannabinoids have been reported to be upregulated in neurodegenerative disorders such as Parkinson’s and Alzheimer’s diseases and reduce unwanted effects or slow disease progression [41, 42]. Additionally, CB_1_Rs have also been reported to be upregulated in the CNS of PLWH [43] and in simian immunodeficiency virus (SIV) encephalitis [44].

Although CB_1_R agonists are effective therapeutic targets in neurodegenerative and neuroinflammatory diseases [45, 46], their cannabimimetic side-effects, including abuse liability and dependence, limit their therapeutic use [47, 48, 49]. CB_1_R agonists such as Δ^9^-tetrahydrocannabinol (THC) and CP55940 directly bind to the orthosteric binding pocket and initiate global activation of the receptor; therefore, it is difficult to isolate the desired effects from the side-effects by direct CB_1_R activation. One approach, to reduce the unwanted side-effects, is by indirectly modulating the activity of CB_1_Rs using allosteric modulators. Allosteric modulators bind to topographically distant sites from the orthosteric site on the receptor. They do not activate the receptor directly but induce a conformational change by binding to an allosteric site that alters ligand potency and/or efficacy [50, 51, 52, 53]. It is hypothesized that CB_1_R-positive allosteric modulators (PAMs) would show neuroprotective effects in the context of HAND and bypass the cannabimimetic side effects [53, 54, 55, 56].

ZCZ011 belongs to the 2-phenylindole class of compounds and is one of the first CB_1_R positive allosteric modulators reported in the literature [55, 57]. Studies have reported ZCZ011 as an ago-PAM, which defines a PAM with intrinsic efficacy in the absence of orthosteric ligand [55]. *In vitro* data show that ZCZ011 increased the binding of CB_1_R agonist [^3^H]CP55940 and potentiated AEA signaling [55]. Additionally, ZCZ011 alone was found to be an agonist in the cAMP assay and a weak agonist in β-arresting recruitment assay [55]. On the contrary, a recent study showed ZCZ011 to behave as an allosteric agonist with an *in vitro* signaling profile similar to THC [57]. In *in vivo* studies, ZCZ011 alone significantly reversed mechanical and cold allodynia in neuropathic pain model without any cannabimimetic effects [55]. Although CB_1_R PAMs show protective effects in various neuropathologies [55, 56, 58, 59, 60], its use as a potential therapeutic target for HAND is unknown.

In the present study, we used HIV-1 Tat transgenic mice to evaluate chronic (14 days) effects of racemic ZCZ011 (10 mg/kg) on various behavioral paradigms, including pain sensitivity, motor activity, motor coordination, anxiety, and object recognition memory. We further investigated the effects of chronic ZCZ011 treatment on the endocannabinoid system in different CNS regions, including the prefrontal cortex, striatum, cerebellum, and spinal cord, by quantifying endogenous cannabinoid and cannabinoid-like ligand concentration via ultraperformance liquid chromatography/tandem mass spectrometry (UPLC-MS/MS). Western blot analysis was conducted to quantify cannabinoid receptors, and endocannabinoid degradative enzymes in various CNS regions (**Fig 1**). Here, we demonstrate the first evidence for therapeutic potential of CB_1_R PAM in the context of neuroHIV.

**Fig 1.**
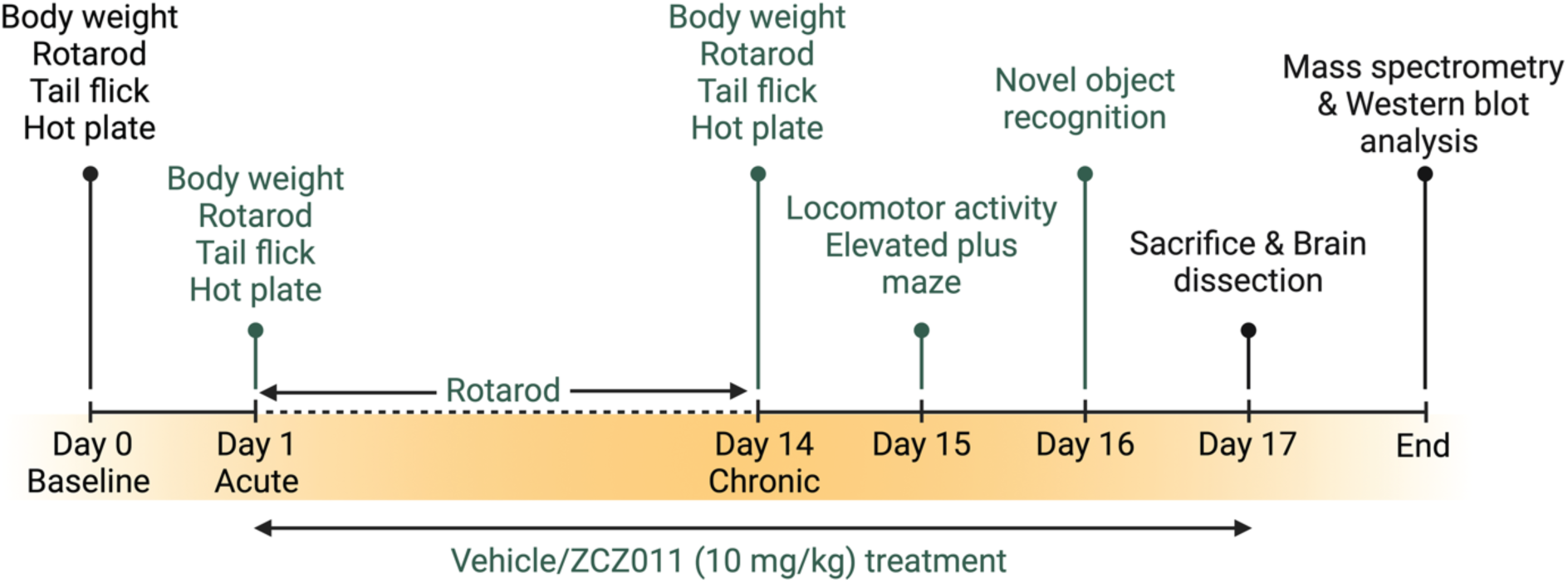
Schematic representation of the experimental study design. Both Tat(–) and Tat(+) mice received DOX-containing chow for 2 months and were trained on the rotarod for 3 days. Subsequently, a baseline (Day 0) recording for body weight, rotarod, tail flick, and hot plate was conducted. Next, vehicle and ZCZ011 (10 mg/kg) were administered for 14 days, and body weight, tail flick, and hot plate were repeated on Day 1 and Day 14 to assess acute and chronic effects, respectively. Rotarod trials were continued throughout drug treatment (Day 1 – Day 14). Animals were habituated to the locomotor activity chamber on Day 14 and tested for locomotor activity and elevated plus maze 60 min after vehicle or ZCZ011 treatment on Day 15. On day 16, animals underwent the training phase for the novel object recognition task. After the training phase, testing was conducted 60 minutes following vehicle or ZCZ011 injection treatment. Finally, on day 17, the animals received their last vehicle or ZCZ011 treatment and were sacrificed after 60 min. Brains were dissected and snap frozen, and stored at −80°C. The right and left hemispheres of the brains were used for mass spectrometry and western blot analysis, respectively. Figure created with BioRender.com.

## Results

### Body mass

Body mass (g) of all animals was taken to ensure the experimental design did not affect animal’s health. Animals were weighed at baseline (Day 0, **Fig 2A**) and after acute (Day 1) and chronic (Day 14) vehicle or ZCZ011 (10 mg/kg) treatment (**Fig 2B**). A two-way ANOVA for baseline and a four-way mixed ANOVA for acute and chronic treatment did not show any significant effects or interactions.

**Fig 2.**
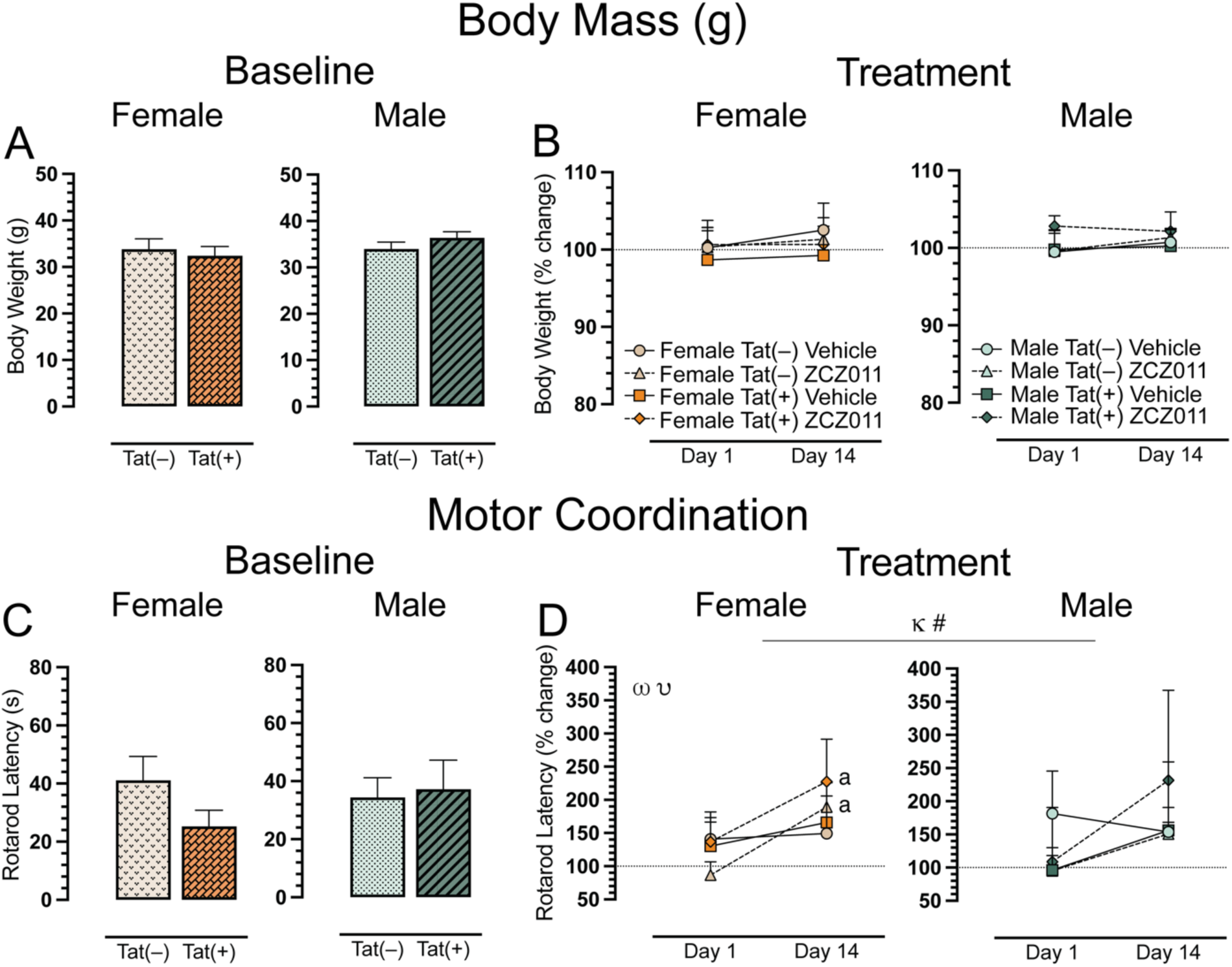
Effect of acute and chronic ZCZ011 treatment (10 mg/kg) on body weight and motor coordination. (**A**) Baseline body weight (g) of female and male mice was taken before starting drug treatment. No significant differences were found in body weight for mice based on sex or genotype. (**B**) Body weight is represented as percent change from baseline, set at 100%, after acute (Day 1) and chronic (Day 14) treatment. No significant differences were noted for either females and males. (**C**) No significant differences were found in the rotarod latency (s) at baseline for females and males. (**D**) Significant effects were noted for rotarod latency, represented as percent change from baseline, set at 100%, after acute and chronic vehicle or ZCZ011 treatment. All data are expressed as mean ± the standard error of the mean (SEM). Statistical significance was assessed by overall ANOVAs; ^κ^*p* = 0.01 main effect of time; non-significant trend, ^#^*p* = 0.08 main time x drug interaction. Separate ANOVAs for females and males; ^ω^*p* = 0.01 main effect of time for females; ^υ^*p* = 0.05 main drug x time interaction for females. Follow-up Tukey’s post hoc tests; ^a^*p* < 0.05 ZCZ011 treated Tat(–) and Tat(+) female mice on Day 1 vs. Day 14. ZCZ011 dose = 10 mg/kg. *N* = 32(16f).

### Motor coordination

The rotarod test was conducted to investigate effects of ZCZ011 treatment (60 min following injection) on motor coordination and function. Baseline assessment revealed no significant effects for rotarod latency (**Fig 2C**). For acute and chronic treatment (**Fig 2D**), a four-way mixed ANOVA revealed a significant main effect of time, *F*(1, 24) = 6.6, *p* = 0.01, with latency to stay on the rotarod increasing over time (Day 1 vs. Day 14). A non-significant trend for time x drug interaction was noted, *F*(1,24) = 3.1, *p* = 0.08, with drug treatment trending to show differential effects on rotarod latency based on acute (Day 1) compared to chronic (Day 14) treatment. Separate ANOVAs for sex, demonstrated for females, a main effect of time, *F*(1,24) = 6.6, *p* = 0.01, with an increase in rotarod latency over time, that was significantly altered by drug, drug x time interaction, *F*(1,12) = 4.5, *p*

= 0.05. Follow-up Tukey’s post hoc tests demonstrated that chronic ZCZ011 treatment significantly increased rotarod latency compared to acute ZCZ011 treatment (*p* = 0.002). This increase in rotarod latency for females was true for both genotypes, Tat(–) females (*p* = 0.01) and Tat(+) females (*p* = 0.02). No significant effects were seen for male mice. One sample *t*-tests did not reveal any significant differences from baseline for females or males.

### Spontaneous heat-evoked nociception

The tail-flick and hot-plate assays were conducted for baseline, acute, and chronic ZCZ011 treatment after rotarod performance, to evaluate heat-evoked pain-like behaviors. For the spinal-related tail-flick assay, a two-way ANOVA demonstrated no significant effect at baseline (**Fig 3A**). For acute and chronic treatment, a four-way mixed ANOVA revealed a significant main effect of time, *F*(1, 24) = 6.7, *p* = 0.01, where tail-flick latency was increased following chronic treatment (Day 14) compared to acute treatment (Day 1). Interestingly, one sample *t*-tests revealed that acute ZCZ011-treated Tat(+) males demonstrated hypersensitivity compared to baseline, *t*(3) = −9.5, *p* = 0.008 (**Fig 3B**).

**Fig 3.**
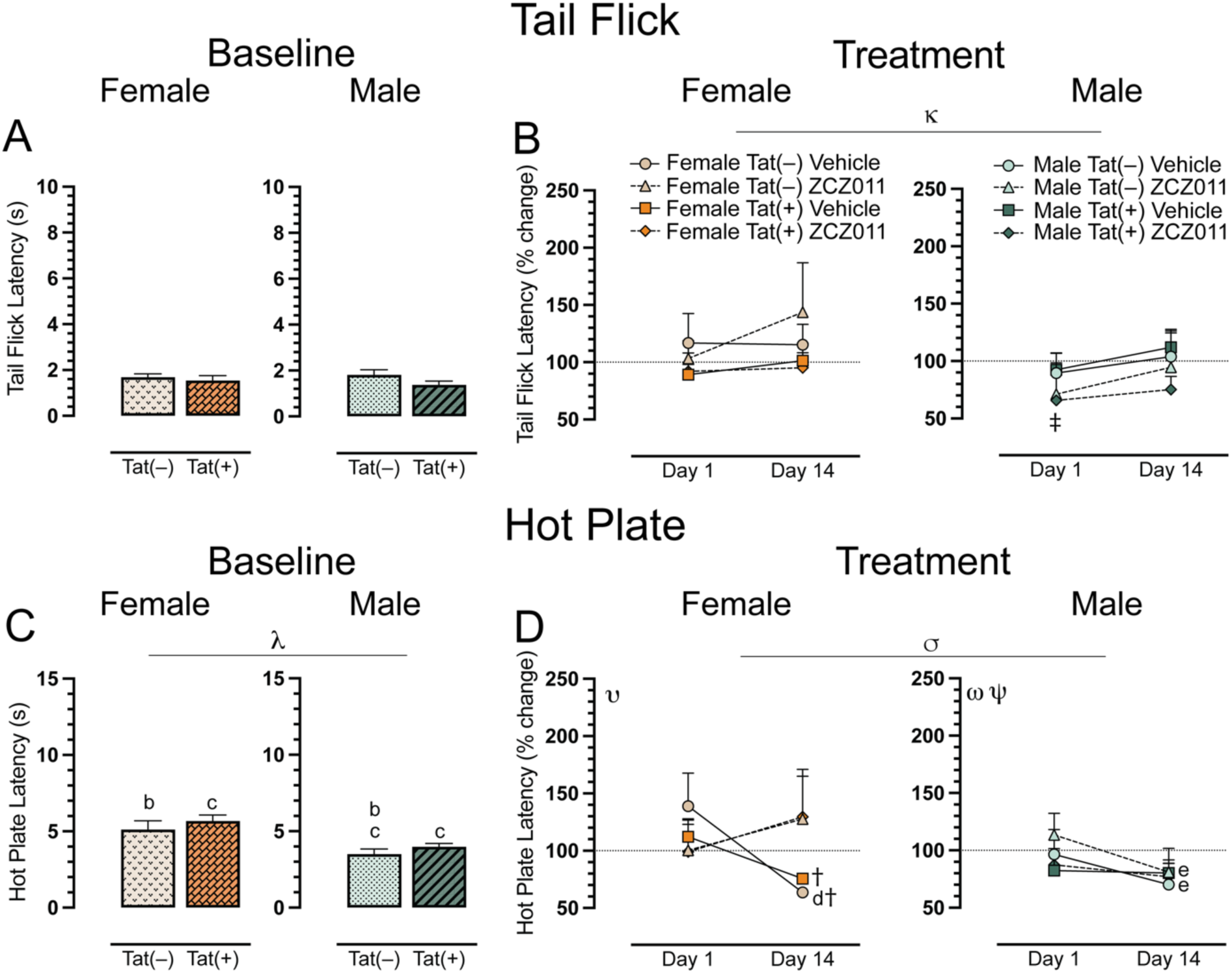
Effect of acute and chronic ZCZ011 treatment (10 mg/kg) on spontaneous evoked nociception. (**A**) Baseline tail flick latency (s) for female and male mice before starting drug treatment. No significant differences were found in the tail flick latency for mice based on sex or genotype. (**B**) Tail flick latency is represented as percent change from baseline, set at 100%, after acute and chronic vehicle or ZCZ011 treatment. A main effect of time was noted after chronic ZCZ011 treatment. (**C**) For baseline hot plate latency (s) females showed higher latency to withdraw paws as compared to males. (**D**) Significant effects were noted for hot plate latency, represented as percent change from baseline, set at 100%, after acute and chronic vehicle or ZCZ011 treatment. All data are expressed as mean ± the standard error of the mean (SEM). Statistical significance was assessed by overall ANOVAs; ^κ^*p* = 0.01 main effect of time; ^λ^*p* <0.001 main effect of sex; ^σ^*p* = 0.008 main time x drug interaction. Separate ANOVAs for females and males; ^υ^*p* = 0.006 time x drug interaction for females; ^ω^*p* = 0.005 main time effect for males; ^ψ^*p* =0.04 time x genotype interaction for males. Follow-up Tukey’s post hoc tests; ^a^*p* = 0.04 baseline Tat(–) males vs baseline Tat(–) females; ^c^*p* < 0.05 baseline Tat(+) females vs baseline Tat(–) and Tat(+) males; ^d^*p* = 0.02 acute vs. chronic vehicle-treated females on day 14; ^e^*p* < 0.05 vehicle and ZCZ011-treated Tat(–) males on Day 1 vs. Day 14. One-sample *t*-test; ^‡^*p* <0.05 ZCZ011-treated Tat(+) males vs. baseline (100%), ^†^*p* <0.05 vehicle-treated Tat(–) and Tat(+) females vs. baseline (100%). ZCZ011 dose = 10 mg/kg. *N* = 32(16f).

The hot-plate assay was used to assess supraspinal-related spontaneous nociception. A two-way ANOVA for the baseline hot-plate latency revealed a significant main effect of sex, *F*(1, 28) = 6.7, *p* < 0.001, with female mice showing higher latencies to withdrawal or lick their paws (**Fig 3C**). A follow up Tukey’s post hoc test revealed significant differences in hot-plate latency based on sex and genotype with Tat(–) females showing higher latency as compared to Tat(–) males (*p* = 0.04), and Tat(+) females displaying higher hot-plate latency as compared to both Tat(–) (*p* = 0.004) and Tat(+) (*p* = 0.03) males. For acute and chronic treatment, a four-way mixed ANOVA indicated a significant time x drug interaction, *F*(1,24) = 8.3, *p* = 0.008, with treatment decreasing hot plate latency over time (**Fig 3D**). Separate three-way mixed ANOVAs for each sex demonstrated for females a significant time x drug interaction, *F*(1,12) = 10.9, *p* = 0.006. Follow-up Tukey’s post hoc tests indicated significant differences between acute and chronic vehicle-treated female mice, specifically found in Tat(–) female mice (*p* = 0.02), with significantly increased hypersensitivity in the hot-plate assay on Day 14. Moreover, the latency for chronically vehicle-treated Tat(–) and Tat(+) females significantly reduced as compared to baseline, *t*(3) = −13.0, *p* = 0.002, and, *t*(3) = −5.4, *p* = 0.048, respectively. For males, we found a main effect of time, *F*(1,12) = 12.1, *p* = 0.005, and a time x genotype interaction, *F*(1,12) = 5.1, *p* = 0.04, with hot plate latency decreasing at Day 14. Tukey’s post hoc test revealed that Tat(–) males showed a significant difference between acute and chronic treatment, for both vehicle (*p* = 0.02) and ZCZ011 (*p* = 0.008) treatments. A significant reduction in hot-plate latency was noted in Tat(–) male mice after chronic treatment with either vehicle or ZCZ011. This effect was not seen in Tat(+) male mice. One sample *t*-tests did not reveal any significant differences from baseline for males.

### Locomotor activity

To evaluate the effects of chronic ZCZ011 exposure on overall activity in Tat transgenic mice, we assessed locomotor activity 60 min after injection. A three-way ANOVA demonstrated no significant effects for locomotor activity after chronic ZCZ011 treatment (**Fig 4A**).

**Fig 4.**
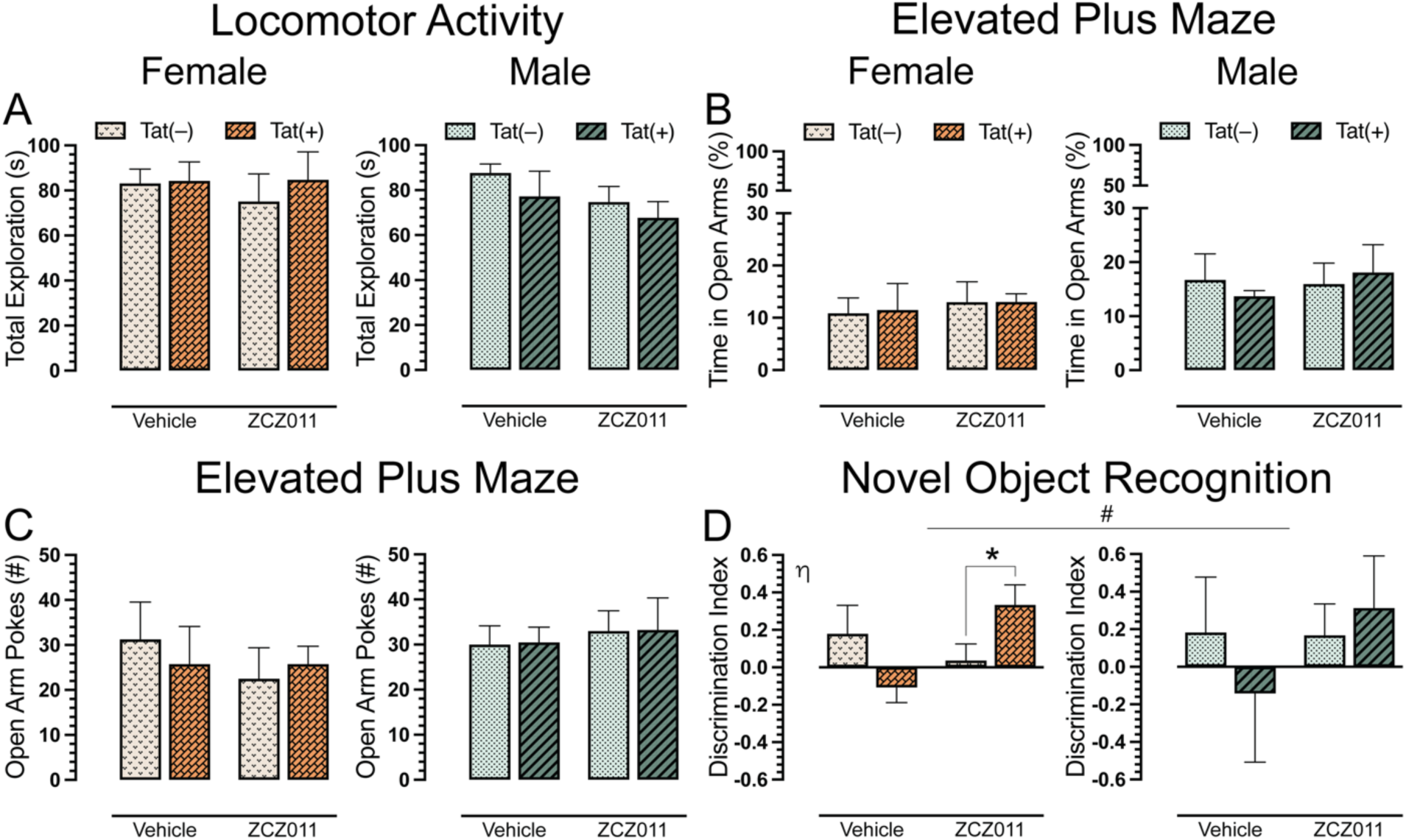
Effect of chronic ZCZ011 treatment on locomotor activity, elevated plus maze, and novel object recognition tasks. (**A**) No significant effects or interactions were noted for total exploration following chronic vehicle or ZCZ011 treatment. (**B**) Percent time spent in open arm during the elevated plus maze was not altered by sex, genotype, or treatment. (**C**) No significant effects were noted for the number of pokes into the open arm. (**D**) Novel object recognition memory is indicated by the discrimination index (0 = no preference, 1 = complete preference for the novel object, −1 = complete preference for the familiar object). A significant improvement in recognition memory was seen in Tat(+) females following ZCZ011 treatment. No changes in recognition memory was observed in male mice. All data are expressed as mean ± the standard error of the mean (SEM). Statistical significance was assessed by overall ANOVAs; non-significant trend, ^#^*p* = 0.09 main drug x genotype interaction. Separate ANOVAs for females and males; ^η^*p* = 0.02 drug x genotype interaction for females. Follow-up Tukey’s post hoc tests; \**p* = 0.01 ZCZ011-treated Tat(–) vs. Tat(+) females. ZCZ011 dose = 10 mg/kg. *N* = 32(16f).

### Anxiety-like behavior

Following the locomotor activity assessment, Tat transgenic mice were tested in elevated plus maze task for anxiety-like behavior. Dependent measures included the percentage of time spent in open arms and the number of pokes into open arms. For percent time spent in open arms (**Fig 4B**), a three-way ANOVA did not show any significant sex, genotype or treatment effect. Similarly, for number of pokes (**Fig 4C**), no significant effects were noted.

### Novel object recognition

A novel object recognition (NOR) task was performed to access recognition memory in Tat transgenic mice following chronic ZCZ011 treatment. Discrimination index was used to measure the animal’s preference between a familiar and novel object, with 1 indicating a complete preference for the novel object, 0 equals no preference, and −1 indicating a complete preference for the familiar object. The three-way ANOVA, revealed a non-significant trend for drug x genotype interaction, *F*(1,24) = 2.9, *p* = 0.09, with ZCZ011 trending to differentially affect recognition memory in Tat(–) and Tat(+) mice. Based on the trend and previously published data [61], a separate three-way ANOVA for sex was conducted. For females, a significant drug x genotype interaction was observed, *F*(1,12) = 6.9, *p* = 0.02, with ZCZ011 treatment differentially affecting recognition memory based on genotype (**Fig 4D**). Follow-up Tukey’s post hoc tests showed that ZCZ011 significantly improved recognition memory in Tat(+) female mice compared to Tat(–) females (*p* = 0.01). No significant effects were observed for male mice.

### CNS levels of endocannabinoids and related lipids

To assess the impact of chronic ZCZ011 (10 mg/kg) exposure on the endocannabinoid system, alterations in levels of AEA, 2-AG, PEA, OEA, and AA were assessed following behavioral experiments in CNS regions, including prefrontal cortex, striatum, cerebellum, and spinal cord of Tat transgenic female and male mice. **Fig 5** shows data from prefrontal cortex, striatum, and cerebellum only, all other data can be found in **S1_Table**. One-way repeated ANOVAs for each lipid revealed that lipid molecule concentration (nmol/g) differed significantly between CNS regions; AEA, *F*(3,93) = 29.3, *p* <0.001, demonstrated differences in expression levels between prefrontal cortex, cerebellum, and spinal cord (*p*’s <0.001), with highest levels of AEA being found in the prefrontal cortex followed by striatum and cerebellum, and the lowest found in the spinal cord. 2-AG, *F*(3,93) = 54.3, *p* <0.001, was found to be highest in the spinal cord followed by cerebellum, striatum and prefrontal cortex (*p*’s <0.001). PEA, *F*(3,93) = 23.5, *p* <0.001, was found to be higher in the spinal cord as compared to cerebellum and spinal cord (*p*’s <0.001). OEA, *F*(3,93) = 10.8, *p* <0.001, was highest in the striatum, followed by spinal cord, cerebellum, and prefrontal cortex (*p*’s <0.001). Lastly, AA, *F*(3,93) = 8.2, *p* <0.001, showed highest levels in cerebellum, followed by striatum, prefrontal cortex, and spinal cord (*p*’s <0.001).

**Fig 5.**
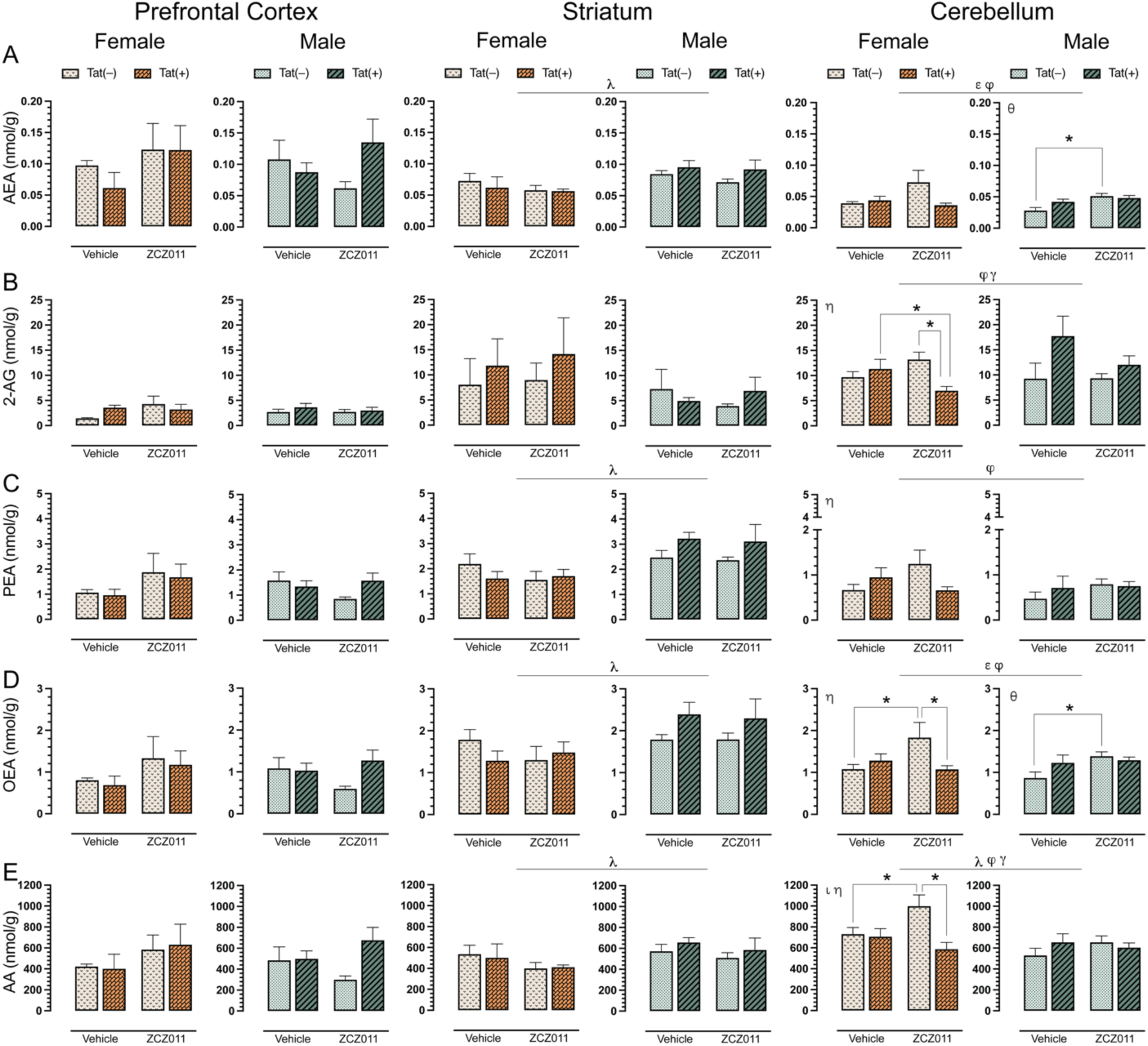
Effect of chronic ZCZ011 treatment on endocannabinoids and related lipids in various CNS regions. Concentration (nmol/g) of (**A**) AEA, (**B**) 2-AG, (**C**) PEA, (**D**) OEA, and (**E**) AA were assessed in the prefrontal cortex, striatum, and cerebellum for vehicle and ZCZ011 treated Tat(–) and Tat(+) mice using LC/MS/MS. In the prefrontal cortex, none of the endocannabinoids and related lipids displayed any main or interaction effects. In the striatum, a main effect of sex was noted, with males showing higher levels of AEA, PEA, OEA, and AA as compared to females. In the cerebellum, chronic ZCZ011 treatment differentially affected the levels of endocannabinoids and related lipids based on sex and/or genotype. All data are expressed as mean ± the standard error of the mean (SEM). Statistical significance was assessed by overall ANOVAs; ^λ^*p* ≤ 0.05 main effect of sex; ^ε^*p* ≤ 0.005 main effect of drug; ^φ^*p* ≤ 0.05 main drug x genotype interaction; ^γ^*p* ≤ 0.05 main drug x sex interaction. Separate ANOVAs for females and males; ^θ^*p* ≤ 0.05 drug effect for males; ^ι^*p* ≤ 0.05 genotype effect for females; ^η^*p* ≤ 0.05 drug x genotype interaction for females. Follow-up Tukey’s post hoc tests; \**p* ≤ 0.05. ZCZ011 dose = 10 mg/kg. *N* = 32(16f).

For AEA (**Fig 5A**), no significant effects were noted in the prefrontal cortex. In the striatum, a three-way ANOVA revealed a main effect of sex, *F*(1,24) = 9.9, *p* = 0.004, with male mice displaying higher levels of AEA as compared to females. No other interactions were observed in the striatum. Additionally, a separate two-way ANOVA for each sex did not show any significant effects in AEA levels for female or male mice. In the cerebellum, a main effect of drug, *F*(1,24) = 6.0, *p* = 0.02, was noted with ZCZ011 treatment increasing AEA levels as compared to vehicle. Further, a drug x genotype interaction, *F*(1,24) = 6.9, *p* = 0.01, was noted, with ZCZ011 treatment differentially affecting AEA levels in Tat(–) and Tat(+) mice. A separate two-way ANOVA for each sex did not show any significant effects in AEA levels for female mice; however, for males a significant effect of drug, *F*(1,12) = 11.0, *p* = 0.006, was noted, with ZCZ011 treatment increasing AEA levels in male mice. A follow-up Tukey’s post hoc test revealed that AEA levels were increased in ZCZ011-treated Tat(–) males as compared to vehicle-treated Tat(–) males (*p* = 0.003).

For 2-AG (**Fig 5B**), both prefrontal cortex and striatum, did not show any significant effects. In the cerebellum, a drug x genotype interaction, *F*(1,24) = 4.9, *p* = 0.03, and a genotype x sex interaction, *F*(1,24) = 6.6, *p* = 0.01, were noted, with Tat expression differentially affecting 2-AG levels based on treatment and sex. A separate two-way ANOVA for each sex revealed a drug x genotype interaction in female mice, *F*(1,12) = 7.7, *p* = 0.01, with ZCZ011 treatment decreasing 2-AG levels as compared to vehicle. A follow-up Tukey’s post hoc test indicated that ZCZ011 treatment decreased 2-AG levels in Tat(+) females as compared to vehicle-treated Tat(+) females (*p* = 0.05) and ZCZ011-treated Tat(–) females (*p* = 0.009). Male mice did not show any main or interaction effects. For PEA (**Fig 5C**), no significant effects were noted in the prefrontal cortex. In the striatum, a main effect of sex was noted, *F*(1,24) = 16.0, *p* <0.001, with male mice displaying higher levels of PEA as compared to females. A separate two-way ANOVA for each sex did not show any significant effects for female or male mice. In the cerebellum, a drug x genotype interaction was observed, *F*(1,24) = 4.8, *p* = 0.03, with ZCZ011 treatment differentially affecting PEA levels in Tat(–) and Tat(+) mice. A two-way ANOVA for each sex indicated a drug x genotype interaction for females, *F*(1,12) = 4.6, *p* = 0.05, with ZCZ011 treatment increasing PEA levels in Tat(–) females and decreasing levels in Tat(+) females. However, follow-up Tukey’s post hoc test did not show any significant differences. Additionally, no significant effects were observed for male mice.

For OEA (**Fig 5D**), no significant changes were seen in the prefrontal cortex. In the striatum, a main effect of sex was noted, *F*(1,24) = 9.5, *p* <0.001, with male mice displaying higher levels of PEA as compared to females. A separate two-way ANOVA for each sex did not show any significant effects in OEA levels for female and male mice. In the cerebellum, a main effect of drug, *F*(1,24) = 5.0, *p* = 0.03, and drug x genotype interaction, *F*(1,24) = 8.0, *p* = 0.009, were noted, with ZCZ011 treatment differentially affecting OEA levels for Tat(–) and Tat(+) mice. A separate two-way ANOVA for each sex showed a significant drug x genotype interaction *F*(1,12) = 5.3, *p* = 0.04 in female mice. A follow-up Tukey’s post hoc test indicated that ZCZ011 treatment increased OEA levels in Tat(–) females as compared to vehicle-treated Tat(–) females (*p* = 0.01) and decreased OEA levels in Tat(+) females as compared to ZCZ011-treated Tat(–) females (*p* = 0.02). In males, a significant drug effect was seen, *F*(1,12) = 4.6, *p* = 0.05, with ZCZ011 treatment increasing OEA levels as compared to vehicle. Follow-up Tukey’s post hoc test showed that chronic ZCZ011 treatment increased OEA levels in Tat(–) males as compared to vehicle-treated Tat(–) males (*p* = 0.01), but not in Tat(+) males.

For AA (**Fig 5E**), no significant effects were observed in the prefrontal cortex. In the striatum, a main effect of sex was noted, *F*(1,24) = 4.3, *p* <0.001, with male mice exhibiting higher levels of AA as compared to females. A separate two-way ANOVA for each sex did not show any significant effects in PEA levels for female and male mice. In the cerebellum, a main effect of sex was observed, *F*(1,24) = 7.7, *p* = 0.01, with females showing higher levels of AA as compared to males. Moreover, a drug x genotype interaction, *F*(1,24) = 7.3, *p* = 0.01 and a genotype x sex interaction, *F*(1,24) = 6.0, *p* = 0.02, were also seen, with Tat expression differentially affecting AA levels based on treatment and sex. A separate two-way ANOVA for each sex indicated, a genotype effect, *F*(1,12) = 7.3, *p* = 0.01, and a drug x genotype interaction, *F*(1,12) = 5.7, *p* = 0.03 for females. A follow-up Tukey’s post hoc tests demonstrated that ZCZ011 treatment increased AA levels in Tat(–) females as compared to vehicle-treated Tat(–) females (*p* = 0.03) and decreased AA levels in Tat(+) females as compared to ZCZ011-treated Tat(–) females (*p* = 0.003). No significant effects were seen in male mice.

Overall, chronic ZCZ011 treatment did not change the levels of endocannabinoids and related lipids in the prefrontal cortex. In the striatum, except for 2-AG, males displayed higher levels of AEA, PEA, OEA, and AA levels as compared to females. Lastly, ZCZ011 had differential effects in the cerebellum where the levels of endocannabinoids and related lipids were dependent on sex and genotype.

### CNS expression levels of cannabinoid receptors, CB_1_R and CB_2_R

To evaluate the effect of chronic ZCZ011 (10 mg/kg) exposure on the endocannabinoid system, alterations in levels of cannabinoid receptors, CB_1_R and CB_2_R, were assessed in CNS regions, including prefrontal cortex, striatum, hippocampus, cortex, cerebellum, brainstem, and spinal cord of Tat transgenic female and male mice. **Fig 6** shows data from the prefrontal cortex, striatum and cerebellum only, due to their relevance to the behavioral studies, the data for other CNS regions can be found in **S2_ Table**. Representative bands from each group is shown in **Fig. 6A** and raw and unedited blots can be found in **Supporting information**. One-way repeated ANOVAs for CB_1_R, *F*(6,186) = 9.1, *p* < 0.001, and CB_2_R, *F*(6,186) = 3.4, *p* = 0.003, demonstrated significant differences between various CNS regions. The order of CB_1_R expression from highest to lowest is as follows, prefrontal cortex > brainstem > spinal cord > hippocampus > striatum > cerebellum > cortex. Similarly, the order of CB_2_R expression from highest to lowest is, cortex > prefrontal cortex > brainstem > cerebellum > striatum > hippocampus > spinal cord.

**Fig 6.**
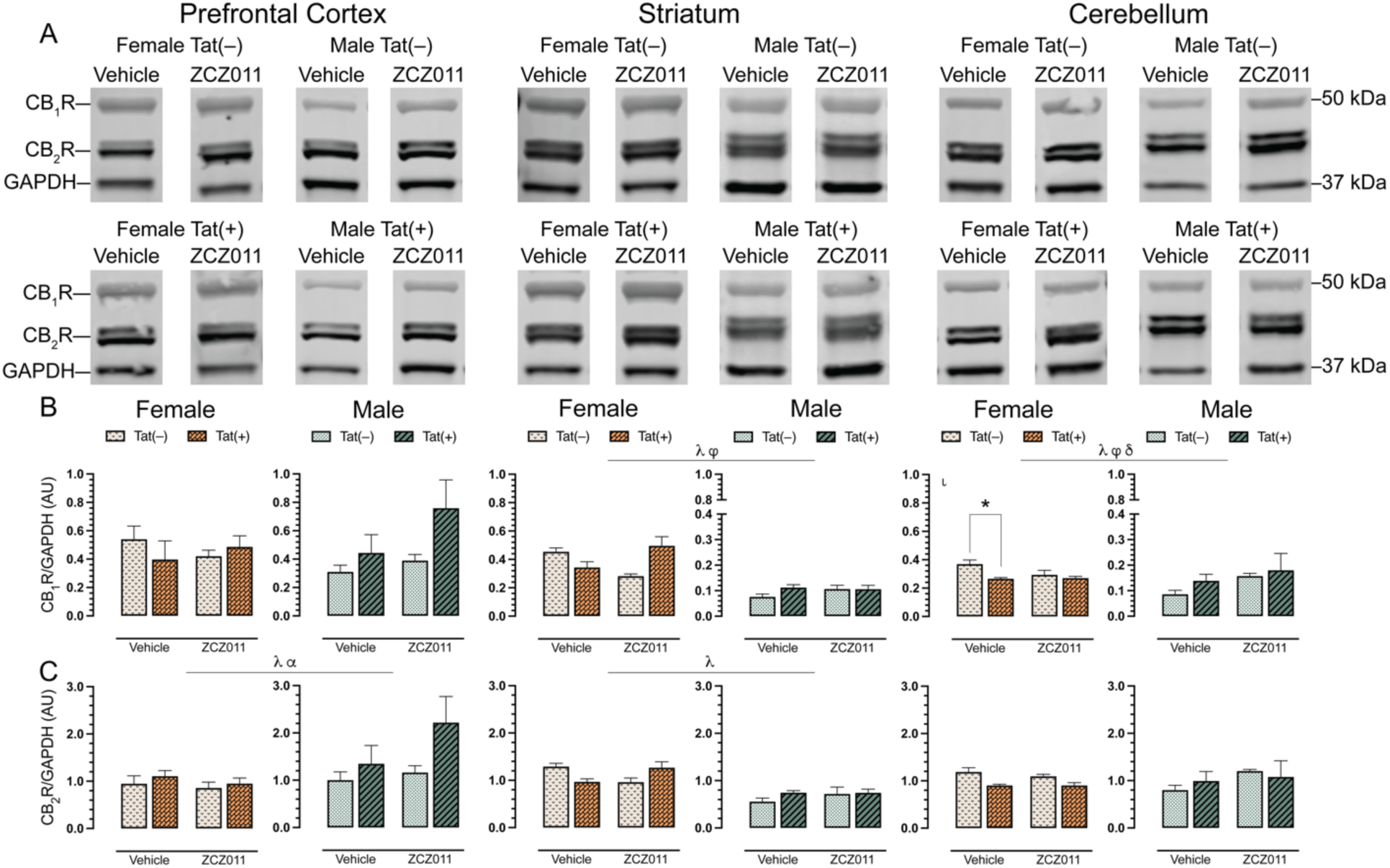
Effect of chronic ZCZ011 treatment on CB_1_R and CB_2_R expression in various CNS regions. CB_1_R and CB_2_R expression levels were assessed in the prefrontal cortex, striatum, and cerebellum for vehicle- and ZCZ011-treated Tat(–) and Tat(+) mice via Western blot analysis. Data were normalized to the housekeeping protein GAPDH. (**A**) Representative immunoblots for CB_1_R, CB_2_R, and GAPDH for all groups used in the study. (**B**) In the prefrontal cortex, no changes were observed in CB_1_R expression. In striatum and cerebellum, a main effect of sex was seen with females displaying higher levels of CB_1_R as compared to males. Additionally, ZCZ011 differentially affected CB_1_R expression based on genotype in the striatum and cerebellum. (**C**) In the prefrontal cortex and striatum, a main effect of sex was seen with males displaying higher level of CB_2_R in the prefrontal cortex and females displaying higher level of CB_2_R in the striatum. Further, Tat expression increased CB_2_R in the prefrontal cortex. In the cerebellum, no changes were observed in CB_2_R expression. All data are expressed as mean ± the standard error of the mean (SEM). Statistical significance was assessed by ANOVAs; ^λ^*p* ≤ 0.05 main effect of sex; ^α^*p* ≤ 0.005 main effect of genotype; ^φ^*p* ≤ 0.05 main drug x genotype interaction; ^δ^*p* ≤ 0.05 main genotype x sex interaction. Separate ANOVAs for females and males; ^ι^*p* ≤ 0.05 genotype effect for females. Follow-up Tukey’s post hoc tests; \**p* ≤ 0.05. ZCZ011 dose = 10 mg/kg. *N* = 32(16f); AU, arbitrary unit.

For CB_1_R (**Fig 6B**), no significant effects were observed in the prefrontal cortex. In the striatum, a three-way ANOVA displayed a main effect of sex, *F*(1,24) = 183.0, *p* < 0.001, with females exhibiting higher levels of CB_1_R as compared to males. Additionally, a drug x genotype interaction was seen, F(1,24) = 10.9, p = 0.003, with ZCZ011 treatment differentially affecting CB_1_R levels based on genotype. A separate two-way ANOVA for each sex did not show any significant effects in CB_1_R expression for female or male mice. In the cerebellum, a main effect of sex was observed, F(1,12) = 54.5, p <0.001, with females showing higher CB_1_R expression as compared to males. Moreover, a drug x sex interaction, *F*(1,12) = 4.5, *p* = 0.04, and a genotype x sex interaction, *F*(1,12) = 5.4, *p* = 0.02, were noted, with ZCZ011 treatment and Tat expression differentially affecting CB_1_R expression based on sex. A separate two-way ANOVA for each sex, demonstrated a genotype effect for females, *F*(1,12) = 7.8, *p* = 0.01, with lower CB_1_R expression in Tat(+) females as compared to Tat(–) females. A follow-up Tukey’s post hoc test displayed that CB_1_R expression was reduced in vehicle-treated Tat(+) females (*p* = 0.007) as compared to Tat(–) female mice. No significant changes were observed in CB_1_R expression in male mice.

For CB_2_R (**Fig 6C**), a three-way ANOVA exhibited a main effect of sex, *F*(1,24) = 4.7, *p* = 0.04, with male mice displaying higher levels of CB_2_R as compared to females, and a main effect of genotype, *F*(1,24) = 4.73, *p* = 0.04, with Tat(+) mice showing higher levels of CB_2_R compared to Tat(–) mice. A separate three-way ANOVAs for each sex did not indicate any significant effects for either females or males. For the striatum, a main effect of sex was observed, *F*(1,12) = 44.8, *p* <0.001, with females displaying significantly higher levels of CB_2_R as compared to males. A separate two-way ANOVA for each sex did not indicate any significant effects. In the cerebellum, CB_2_R expression was not significantly altered.

In summary, CB_1_R and CB_2_R expression was not altered in the prefrontal cortex and cerebellum, respectively. Further, CB_1_R (in the striatum and cerebellum) and CB_2_R (in the prefrontal cortex) expression was altered based on sex, genotype, and treatment.

### CNS expression levels of endocannabinoid degradative enzymes, FAAH and MAGL

To assess the effect of chronic ZCZ011 (10 mg/kg) exposure on the endocannabinoid system, changes in levels of endocannabinoid degradative enzymes, FAAH and MAGL, were evaluated in CNS regions, including prefrontal cortex, striatum, hippocampus, cortex, cerebellum, brainstem, and spinal cord of Tat transgenic female and male mice. **Fig 7** shows data from the prefrontal cortex, striatum and cerebellum only, due to their relevance to the behavioral studies. Data for other CNS regions can be found in **S2_Table**. Representative bands from each group is shown in **Fig 7A** and raw and unedited blots can be found in **Supporting information**. One-way repeated ANOVAs for FAAH, *F*(6,186) = 14.1, *p* < 0.001, and MAGL, *F*(6,186) = 11.3, *p* < 0.001, demonstrated significant differences between various CNS regions. The order of FAAH and MAGL expression from highest to lowest is as follows: FAAH: prefrontal cortex > cortex > hippocampus > cerebellum > spinal cord > striatum > brainstem, and MAGL: cortex > hippocampus > spinal cord > prefrontal cortex > brainstem > cerebellum > striatum.

**Fig 7.**
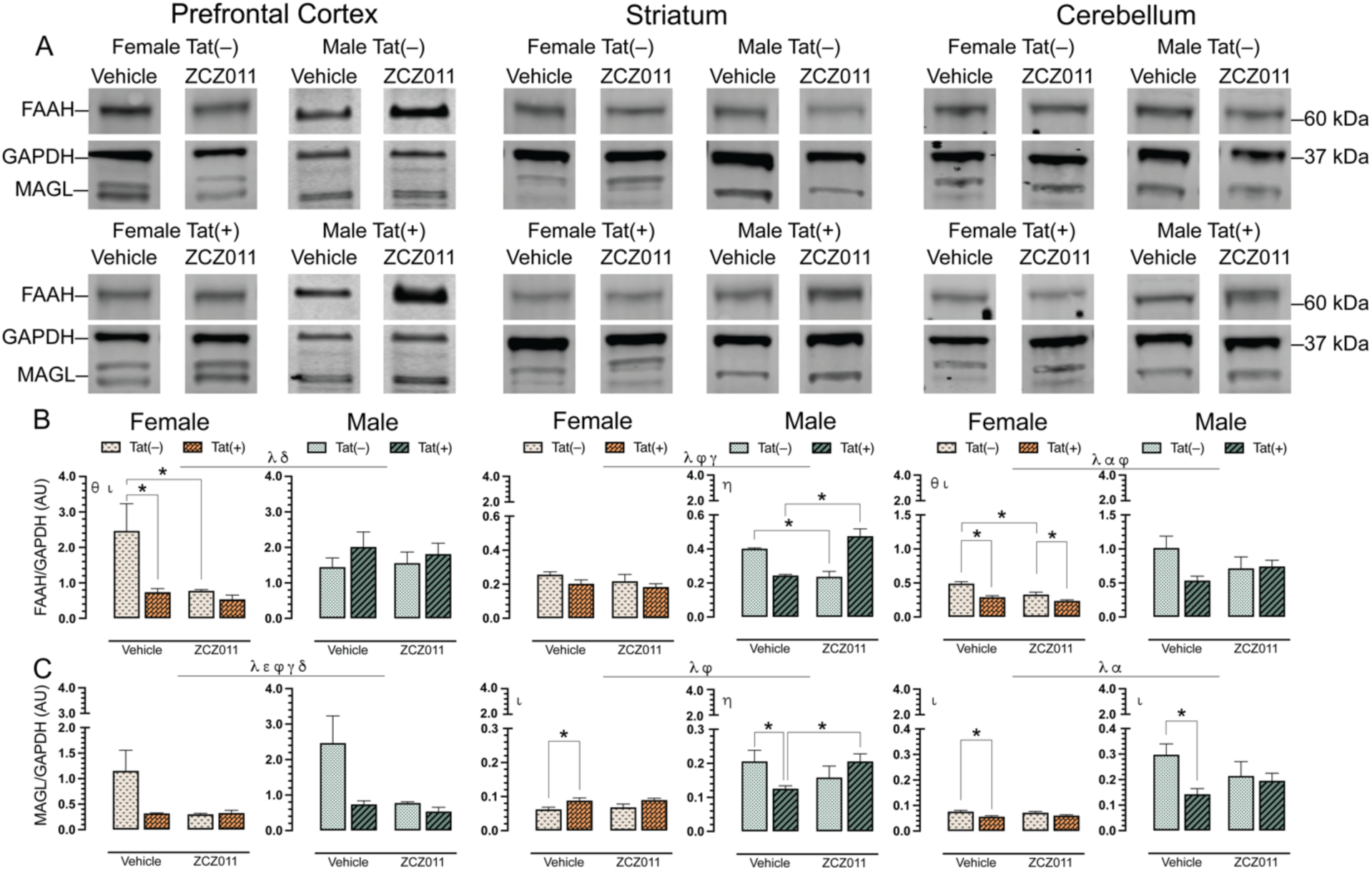
Effect of chronic ZCZ011 treatment on FAAH and MAGL expression in various CNS regions. FAAH and MAGL expression levels were assessed in the prefrontal cortex, striatum, and cerebellum for vehicle- and ZCZ011-treated Tat(–) and Tat(+) mice via Western blot analysis. Data were normalized to the housekeeping protein GAPDH. (**A**) Representative immunoblots for FAAH, MAGL, and GAPDH for all groups used in the study. (**B** and **C**) A main effect of sex was noted in prefrontal cortex, striatum and cerebellum with males displaying higher levels of FAAH and MAGL as compared to females. Additionally, the levels of FAAH and MAGL expression were differentially affected in all the regions based on sex, genotype, and drug treatment. All data are expressed as mean ± the standard error of the mean (SEM). Statistical significance was assessed by ANOVAs; ^λ^*p* ≤ 0.05 main effect of sex; ^α^*p* ≤ 0.005 main effect of genotype; ^ε^*p* ≤ 0.05 main drug effect; ^φ^*p* ≤ 0.05 main drug x genotype interaction; ^γ^*p* ≤ 0.05 main drug x sex interaction; ^δ^*p* ≤ 0.05 main genotype x sex interaction. Separate ANOVAs for females and males; ^θ^*p* ≤ 0.05 drug effect for females; ^ι^*p* ≤ 0.05 genotype effect for females or males; ^η^*p* ≤ 0.05 drug x genotype interaction males. Follow-up Tukey’s post hoc tests; \**p* = ≤ 0.05. ZCZ011 dose = 10 mg/kg. *N* = 32(16f); AU, arbitrary unit.

For FAAH (**Fig 7B**), a three-way ANOVA demonstrated a main effect of sex in the prefrontal cortex, *F*(1,24) = 5.0, *p* = 0.03, with males displaying higher levels as compared to females, and a genotype x sex interaction, *F*(1,24) = 7.4, *p* = 0.01, with FAAH levels being altered based on sex and genotype. A separate two-way ANOVA for each sex indicated a drug effect for females, *F*(1,24) = 5.8, *p* = 0.03, with ZCZ011 treatment decreasing FAAH expression in females as compared to vehicle, and a genotype effect, *F*(1,12) = 6.3, *p* = 0.02, with FAAH expression being lower in Tat(+) females as compared to Tat(–) females. Follow-up Tukey’s post hoc tests demonstrated that FAAH expression was significantly reduced in ZCZ011-treated Tat(–) females (*p* = 0.01) and vehicle-treated Tat(+) females (*p* = 0.009) as compared to vehicle-treated Tat(–) females. No significant effects were noted in males. In the striatum, a main effect of sex was seen, *F*(1,24) = 43.5, *p* <0.001, with males showing higher levels of FAAH expression as compared to females. Moreover, a drug x genotype interaction, *F*(1,24) = 30.39, *p* <0.001, and a genotype x sex interaction, *F*(1,24) = 4.93, *p* = 0.03, was observed, suggesting that Tat expression has differential effects based on ZCZ011 treatment and sex. A separate two-way ANOVA for each sex did not show any significant effects in female mice; however, for male mice, a drug x genotype interaction, *F*(1,12) = 54.07, *p* <0.001, was noted, with ZCZ011 treatment showing differential effects in Tat(–) and Tat(+) males. Follow-up Tukey’s post hoc tests indicated that ZCZ011 treatment reduced FAAH expression in Tat(–) males as compared to vehicle-treated Tat(–) males (*p* <0.001), but increased FAAH expression in Tat(+) males as compared to vehicle-treated Tat(+) males (*p* <0.001). In the cerebellum, a main effect of sex, *F*(1,24) = 37.7, *p* <0.001, and a main effect of genotype, *F*(1,24) = 7.5, *p* = 0.01, were observed, with males and Tat(–) mice displaying higher levels of FAAH expression as compared to females and Tat(+) mice, respectively. Additionally, a drug x genotype interaction was seen, *F*(1,24) = 5.1, *p* = 0.03, with ZCZ011 treatment differentially affecting FAAH expression in Tat(–) and Tat(+) mice. A separate two-way ANOVA for each sex showed a genotype effect in females, *F*(1,12) = 30.8, *p* <0.001, and a drug effect, *F*(1,12) = 16.7, *p* = 0.001, with decreased FAAH expression found in Tat(+) females and upon ZCZ011 treatment, respectively. Follow-up Tukey’s post hoc tests demonstrated that FAAH expression was reduced in both vehicle- (*p* <0.001) and ZCZ011-treated (*p* = 0.02) Tat(+) females as compared to vehicle- and ZCZ011-treated Tat(–) females, respectively. Additionally, ZCZ011-treated Tat(–) mice had significantly lower expression of FAAH (*p* <0.001) as compared to vehicle-treated Tat(–) female mice. Male mice did not show any significant effects.

For MAGL (**Fig 7C**), in the prefrontal cortex a main effect of sex, *F*(1,24) = 23.3, *p* <0.001, and a main effect of drug, *F*(1,24) = 4.1, *p* = 0.05, were observed, with males and vehicle-treated mice showing higher expression of MAGL. Additionally, interactions including drug x genotype, *F*(1,24) = 4.30, *p* = 0.04, drug x sex, *F*(1,24) = 4.23, *p* = 0.05, and genotype x sex, *F*(1,24) = 4.14, *p* = 0.05, were also found. A separate two-way ANOVA for each sex did not indicate any significant effects in either male or female mice. In the striatum, a main effect of sex, *F*(1,24) = 49.9, *p* <0.001, with males showing higher levels of MAGL as compared to females, and a drug x genotype interaction, *F*(1,24) = 5.8, *p* = 0.03, were seen, with ZCZ011 treatment differentially affecting MAGL expression in Tat(–) and Tat(+) mice. A separate two-way ANOVA for each sex indicated a genotype effect in female mice, *F*(1,12) = 9.71, *p* = 0.009, with Tat(+) females demonstrating higher MAGL levels as compared to Tat(–) females. Follow-up Tukey’s post hoc tests indicated that MAGL level were increased in vehicle-treated Tat(+) females as compared to vehicle-treated Tat(–) females (*p* = 0.03). For males, a drug x genotype interaction was observed, *F*(1,12) = 5.91, *p* = 0.03, with ZCZ011 treatment differentially affecting MAGL expression based on genotype. Follow-up Tukey’s post hoc tests indicated that MAGL expression was reduced in vehicle-treated Tat(+) males as compared to vehicle-treated Tat(–) males (*p* = 0.05), and as compared to ZCZ011-treated Tat(+) males (*p* = 0.05). Finally, in the cerebellum, a two-way ANOVA demonstrated a main effect of sex, *F*(1,24) = 54.3, *p* <0.001, and a main effect of genotype, *F*(1,24) = 6.6, *p* = 0.01, with males and Tat(–) expressing significantly higher levels of MAGL as compared to females and Tat(+) mice, respectively. A separate two-way ANOVA for each sex indicated a main genotype effect for both females, *F*(1,12) = 12.6, *p* = 0.003, and males, *F*(1,12) = 4.8, *p* = 0.04 with decreased MAGL expression in Tat(+) mice as compared to Tat(–) mice. Follow-up Tukey’s post hoc tests demonstrated MAGL expression was reduced in vehicle-treated Tat(+) females (*p* = 0.006) and vehicle-treated Tat(+) males (*p* = 0.01) as compared to their respective vehicle-treated Tat(–) counterparts.

Overall, males displayed higher levels of FAAH and MAGL in the prefrontal cortex, striatum, and cerebellum as compared to females. Additionally, ZC011 treatment differentially affected FAAH and MAGL expression based on sex and genotype.

### Relationship between endocannabinoid and related lipids and motor coordination

As the most prominent findings were noted in the cerebellum, we assessed the relationship between FAAH expression and AA levels, in the cerebellum. As no effects were noted in males, correlation studies were only done in females. Pearson correlation were conducted specifically in the cerebellum. Data indicated that in the cerebellum, no correlations were observed in the vehicle-treated females. Moreover, in ZCZ011-treated females a significant positive relationship was noted with higher levels of FAAH expression being associated with higher levels of AA in the cerebellum (**Fig 8A**). Significant correlations were further evaluated by simple regression analysis. Results indicate predictability of AA levels by FAAH expression, *F*(1,6) = 23.6, *p* = 0.002, accounting for 80% in the cerebellum of ZCZ011-treated female mice. Next, we were interested in exploring the relationship between the endocannabinoid and related lipids expression in the cerebellum and the rotarod performance on Day 14 in females. Person correlation did not reveal any correlation in vehicle-treated females; however, in ZCZ011-treated females, a significant positive relationship was observed with higher levels of AEA (**Fig 8B**) and PEA (**Fig 8C**) in the cerebellum being associated with improved rotarod performance. A simple linear regression demonstrated predictability of rotarod performance by AEA and OEA levels. Specifically, in the cerebellum of ZCZ011-treated females, AEA predicted 59% of total variance in rotarod performance data, *F*(1,6) = 8.7, *p* = 0.02 and PEA predicted 49% of total variance in rotarod performance, *F*(1,6) = 5.7, *p* = 0.05.

**Fig 8.**
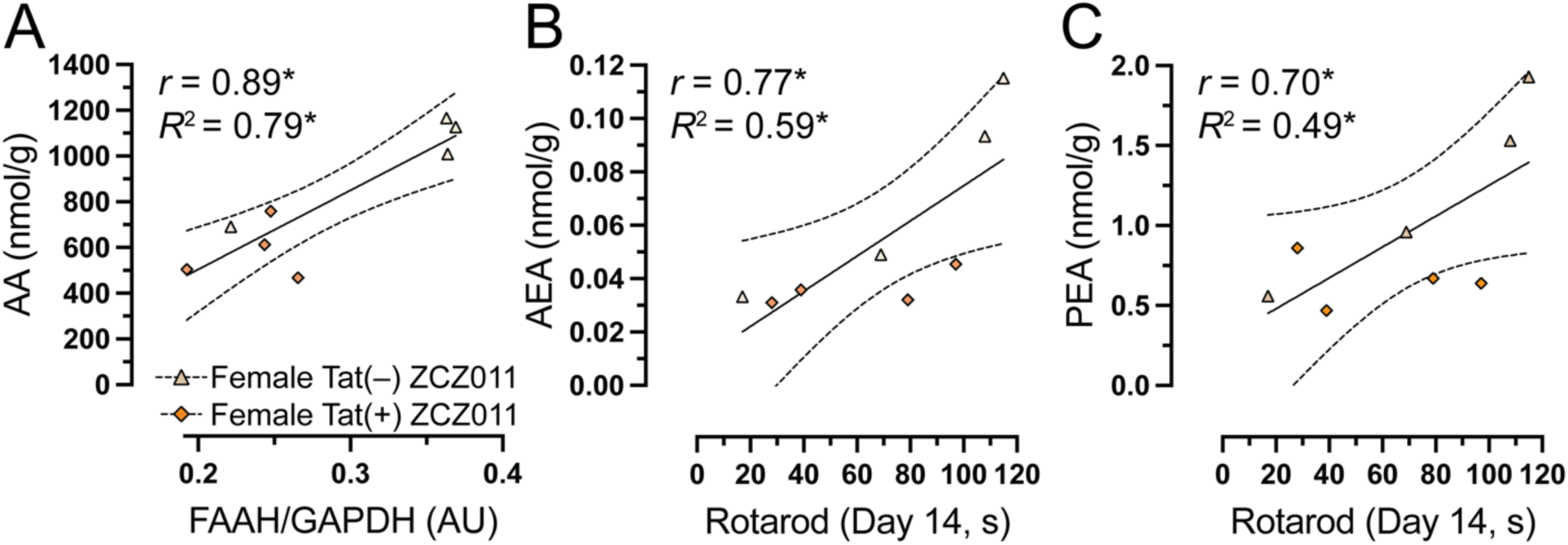
Relationship between endocannabinoid system and motor coordination in female Tat transgenic mice following chronic ZCZ011 treatment. Pearson correlation analysis were followed by simple regression analysis in female Tat(–) and Tat(+) mice to explore the relationship between AA and FAAH expression and between endocannabinoids and related lipids and rotarod performance. (**A**) A significant positive correlation was noted for female with higher AA levels being associated with higher FAAH expression. A significant positive correlation was also seen with higher levels of (**B**) AEA and (**C**) PEA levels being associated with improved rotarod performance on Day 14 in both Tat(–) and Tat(+) female mice. \**p* < 0.05.

## Discussion

This is the first study to explore the effects of CB_1_R PAM in the context of neuroHIV. The unwanted side effects of CB_1_R limit its potential to be targeted for its neuroprotective effects; however, PAMs can bypass these adverse effects and harness the neuroprotective effects of the CB_1_R, making it possible to explore the CB_1_R indirectly. Here, we report the effects of chronic ZCZ011, a CB_1_R PAM, on various behavioral measures and the endocannabinoid system in HIV-1 Tat transgenic mice. Behavioral assessments demonstrated that chronic ZCZ011 treatment enhanced motor coordination and decreased hot plate latency in female mice. Differential effects of chronic ZCZ011 were found for the novel object recognition task upon Tat induction in female mice, with no effects in male mice. Moreover, ZCZ011 treatment altered the endocannabinoid system, dependent on sex, genotype, treatment, and CNS region.

The findings for body mass indicated that chronic ZCZ011 treatment did not affect body mass for either female or male Tat transgenic mice. Studies have shown that ZCZ011 has a CB_1_R-mediated effect in attenuating naloxone-related weight loss [62]. It was observed that in ZCZ011-treated mice, rimonabant (CB_1_R antagonist), but not SR14428 (CB_2_R antagonist), blocked the protective effect of ZCZ011 on naloxone-induced body mass loss. Further, it was also observed that the protective effects of ZCZ011 in attenuating naloxone-related weight loss were lost entirely in CB_1_R knockout mice, which again solidifies that the mechanism of action of ZCZ011 is CB_1_R mediated [62]. Another study showed that in a mouse model for Huntington’s disease CB_1_R PAMs, GAT211, and GAT229 promoted normal weight gain and increased body fat content in R8/2 mice, which is critical, as patients suffering from Huntington’s disease experience a failure to gain weight and body fat content [63]. The current study also found that chronic ZCZ011 treatment improved motor coordination only in female mice. Although no other studies have evaluated CB_1_R PAMs in wildtype mice for motor coordination, in Fisher 344 x Brown Norway F1 rats, a rat model to study age related cognitive deficits, the latency to fall for female mice or rats is higher than that of males [64]. Another study demonstrated that in MAGL knockout and FAAH knockout mice, GAT211 increased immobility time; however, it did not affect motor performance [65]. The increase in rotarod latency for females in the present study may also be due to alterations in the endocannabinoid levels seen in the cerebellum of female mice.

The spontaneous evoked nociception experiments, including tail-flick and hot-plate assays, were interesting as we saw that repeated ZCZ011 treatment increased the tail flick latency in Tat transgenic female and male mice, whereas, for the hot plate, the latency to withdraw or lick their paws was increased only in female mice. It was also observed that the hot-plate latency was reduced in males and vehicle-treated females, possibly due to behavioral tolerance. Behavior tolerance is a learned behavior response that leads to diminished reaction times with repeated measures [66]. Numerous studies in the past have reported a decrease in hot plate latency upon repeated testing [67, 68, 69]; however, it has also been proven that this decrease does not influence the antinociceptive potency of an analgesic compound [70]. Only a handful of studies have evaluated the effects of CB_1_R PAMs with regard to pain. One study has shown that ZCZ011 effectively blocked neuropathic pain and produced anti-allodynic effects in a chronic constriction injury model of neuropathic pain [55]. Additionally, studies using another CB_1_R PAM, GAT211, have been proven to be a very effective and safe analgesic strategy without any dependence and abuse liability [65]. GAT211 exhibited dose-dependent CB_1_R-mediated antinociceptive effects in inflammatory and neuropathic pain models [65]. The current study also demonstrated that the therapeutic efficacy for ZCZ011 was maintained after 14 days of repeated treatment, similar to other CB_1_R PAM studies [55, 65]. Tolerance rapidly develops against CB_1_R agonists [71] and MAGL inhibitors at high doses [72, 73], and the lack of tolerance for CB_1_R PAMs makes them superior targets for antinociceptive properties over traditional targets.

Since PAMs increase the signaling of the endogenous ligand at a receptor, it is essential to evaluate its effect on locomotor activity. A study evaluating GAT211 has shown that, unlike WIN55212, a potent cannabinoid receptor agonist, GAT211 did not alter motor performance, induce immobility, or CB_1_R-dependent withdrawal symptoms in MAGL knockout and FAAH knockout mice [65]. Another study indicated that GAT229 did not produce changes in locomotor activity [74]. Similarly, in the present study, ZCZ011 treatment for over two weeks did not produce any changes in locomotor activity. In the diseased R6/2 mouse model of Huntington’s disease, which shows hypolocomotion, GAT211, GAT228, and GAT229 increased the total distance traveled [63]. Another study has demonstrated that the NMDA antagonist, MK801, produces hyperlocomotion in rats, which was significantly decreased by GAT211 treatment [75]. A potential mechanism by which GAT211 reduced the MK801-induced hyperlocomotion is by inhibiting CB_1_Rs on dysregulated glutamatergic and GABAergic neurons that are involved with modulating motor activity in the cortico-striatal-limbic circuitry responsible for hyperarousal [76, 77]. These data suggest that CB_1_R PAMs can potentially normalize locomotor activity in diseased states, making them an attractive target for treating locomotor activity-related deficits, even though several studies have shown Tat-induced hypolocomotion [78, 79, 80]. The lack of Tat effects on locomotor activity may be related to immune tolerance seen with prolonged Tat exposure [81].

High prevalence of depressive and anxiety disorders in PLWH is supported by past studies [82, 83, 84], in the current study, there was no effect of sex or Tat induction on anxiety-like behavior. However other studies have shown increased anxiety-like behavior in Tat transgenic mouse model [85, 86, 87], which may be due to longer duration of Tat induction (up to 6 months as compared to 2 months in the present study) [85] and at higher doses (via DOX injections as compared to DOX food) [85, 86, 88]. Previous studies from our lab have shown that in the Tat transgenic mouse model, female mice displayed more anxiety than male mice [61]. With regards to CB_1_R PAMs, a previous study in Huntington’s disease mouse model has shown that GAT211, GAT228, and GAT229 increased amount of time mice spent in the central quadrant of the open field test, modeling an anxiolytic effect [63]. On the other hand, GAT211 failed to reduce startle reactivity and prepulse inhibition, a measure of anxiety [89] in MK0801-treated rats [75]. In the present study, no significant differences were seen for the time spent in open arms, and the number of pokes in the open arms, which may be due different neuropathology (Hungtington’s disease vs. HIV-1 Tat) or simply due to different efficacy of CB_1_R PAMs (GAT211 vs ZCZ011). Furthermore, approximately 30-50% of PLWH display memory and learning-related deficits [4, 90, 91]. Prefrontal cortex-related tasks, such as the novel object recognition task, usually show deficits in recognition memory in mice exposed to Tat [92, 93]. In the present study, these effects are only supported in female mice. No effects were seen for males, which may be due to the lack of alterations in the endocannabinoid system or similarities of chosen objects [94, 95]. Additionally chronic ZCZ011 treatment showed differential effects with improving recognition memory only in Tat(+) female mice. ZCZ011 treatment reversed the memory deficits seen in female Tat(+) mice which may be mediated by the endocannabinoid system; however, in the present study no alterations were noted in the endocannabinoids and related lipids in the prefrontal cortex; therefore, additional studies are necessary to decipher the mechanism of action that least to memory related neuroprotective effects.

Alterations in the endocannabinoid system have been reported in the brain of PLWH [43, 96]. Changes in the CB_1_R and CB_2_R expression in the frontal cortex have been reported in postmortem tissue samples of PLWH [43, 96] and FAAH upregulation in cortical postmortem tissue samples of rhesus macaques [44]. Additionally, preclinical studies have also shown Tat-induced changes in endocannabinoid levels and related lipids [97, 98, 99], CB_1_R, CB_2_R, and FAAH expression [61, 100]. However, in the present study we did not see Tat-induced changes in endocannabinoids and related lipids in CNS regions which is similar to a recent study [101] and maybe due to the immune tolerance caused by prolonged Tat exposure [81]. Moreover, not much is known about sex-dependent effects on the endocannabinoid system and its relationship to neuroinflammation and HAND. However, it is known that women show more pronounced HAND symptoms [102, 103, 104, 105] and higher immune activation [106, 107]. In the present study, sex-dpendent alterations were noted for AEA, PEA, OEA, and AA levels in the striatum. Similar to previous study [108], the current study displayed higher AA levels in males compared to females in the prefrontal cortex and striatum, which is not surprising as males also displayed higher levels of AEA, OEA, and PEA, which are metabolized into AA. Moreover, even though CB_1_R and CB_2_R expression was not affected by Tat, females demonstrated higher levels of cannabinoid receptors (CB_1_R and CB_2_R) in prefrontal cortex, striatum, and cerebellum whereas males displayed higher levels of degradative enzymes (FAAH and MAGL). Studies have reported that the endocannabinoid system exhibit sexual dimorphism with females reaching peak CB_1_R levels earlier than males [109, 110], also females showing higher levels of CB_1_ and CB_2_R mRNA levels in all brain regions [111]. Additionally, the levels of estradiol strongly influence the endocannabinoid tone with the receptor numbers and mRNA levels fluctuating across the estrous cycle in females [112, 113, 114]. All these studies suggest that sex differences in the endocannabinoid system develop early and are dependent on circulating hormone levels which may be responsible for the differential effects seen in studies.

Little is known about the effect of CB_1_R PAMs on the endocannabinoid system. A study evaluating mRNA levels of the cannabinoid receptors in Huntington’s disease R6/2 mouse model found that in the diseased model, CB_1_R and CB_2_R mRNA levels were normalized to the wildtype levels in the striatum and cortex of GAT211 treated mice [63]. This suggests that CB_1_R PAMs have the ability to normalize the hyperactivation of the endocannabinoid system in a diseased state. In the present study ZCZ011 treatment displayed a differential effect in the levels of endocannabinoids and related lipids, cannabinoid receptors, and degradative enzymes. Chronic ZCZ011 treatment did not alter CB_1_R or CB_2_R levels in either females or males. Interestingly, ZCZ011 altered the AA levels in the cerebellum of female mice which was positively correlated with FAAH levels in the cerebellum. AEA is metabolized by FAAH into AA and ethanolamine [115], therefore higher levels of FAAH would indicate increased levels of AA which is in line with our findings. Additionally, AEA and PEA levels in females were positively correlated with the rotarod performance suggesting that the improved rotarod performance seen in females may be due to increased levels of AEA in the cerebellum. Studies have suggested that females produce overall higher levels of CB_1_ and CB_2_R mRNA levels as compared to males [111], including cerebellum [116]. Similarly, the present study also indicated that females had higher levels of CB_1_R in the cerebellum as compared to the males and although literature shows that CB_1_R is not expressed in the Purkinje cells in rodents [117], the presence of CB_1_R surrounding area may play a role in regulating the function of the Purkinje cells in the cerebellar cortex [118, 119]. Although we did not see a direct correlation between CB_1_R and the rotarod performance, there is a possibility that the higher levels of CB_1_R causes increase in AEA expression responsible for improving motor coordination which is only observed in females.

There are several limitations to this study which should be taken into consideration when applying these findings to the therapeutic potential of CB_1_R allosteric modulators in the context of HIV. First, is the use of HIV Tat transgenic mouse model. Even though Tat transgenic mouse model is a well-established neuroHIV model, it only expresses one of the many HIV proteins. Therefore, this should be considered when generalizing findings to HIV-1 infection, as different viral proteins may interact and target various signaling pathways in the CNS and behavior in different ways as compared to a single viral protein. Second, using doxycycline to induce Tat expression in our Tat transgenic mouse model is a limitation, as doxycycline has been shown to have neuroprotective effects [120]. In order to control for this confound and minimize bias, both Tat(–) and Tat(+) groups were fed the same doxycycline chow throughout the study. Third, in the present study ZCZ011 was administered alone and subcutaneously as compared to previous studies [55, 62] where intraperitoneal route was used and ZCZ011 was administered along with a CB_1_R orthosteric ligand [55]. These differences in the methodologies may have led to only subtle changes in behavior and the endocannabinoid system following chronic ZCZ011 treatment.

## Conclusion

In conclusion, the present study demonstrated differential effects of chronic ZCZ011 on selected behaviors in a neuroHIV mouse model, which were mediated by sex and HIV Tat expression. Specifically, chronic ZCZ011 treatment improved motor coordination and recognition memory in female mice only. Additionally, chronic Tat expression had some effect on the endocannabinoid system with downregulating CB_1_R, FAAH, and MAGL expression in the cerebellum of female mice. Future studies should be directed towards investigating the role of the estrous cycle and sex hormones that may be responsible for these sex-based effects in the context of HIV. Lastly, as the current study used a racemic mixture, it would be interesting to evaluate the two enantiomers of ZCZ011 to assess if they behave like true PAM and allosteric-agonists similar to GAT211 enantiomers.

## Material and methods

All experiments were approved by the University of North Carolina at Chapel Hill and conducted following the National Institutes of Health Guide for the Care and Use of Laboratory Animals. The method section describes the experimental design used for this study.

### Animals

Doxycycline (DOX)-inducible, brain-restricted HIV-1_IIIB_ Tat_1-86_ transgenic mice were developed on a hybrid C57BL/6J background as described in detail previously [121, 122]. Tetracycline-responsive promoter controlled by glial fibrillary acidic protein (GFAP) expression is responsible for Tat expression, which was induced with a special formulated chow containing 6 mg/g DOX (product TD.09282; Envigo, Indianapolis, IN). At four weeks of age, genotyping was performed to identify the Tat(+) and Tat(−) mice. Inducible Tat(+) transgenic mice express both GFAP-rtTA and TRE-tat genes, whereas control Tat(−) transgenic mice express only the GFAP-rtTA genes. Tat transgenic mice [*n* = 32, (16f); 10 months of age] were held on ad libitum DOX chow for 2 months prior the behavioral experiments and were housed on a reversed 12 h light/dark cycle.

### Drug treatment

For behavioral experiments, animals were chronically injected subcutaneously (s.c.) for 14 days with either vehicle [1:1:18; ethanol, Kolliphor, 0.9% saline] or 10 mg/kg ZCZ011 dissolved in vehicle. The dose of ZCZ011 was selected based on previous study [123]. Administration of vehicle or ZCZ011 was done, 1 h before behavioral tests, via s.c. injections at a volume of 10 mL/g of body mass.

### Experimental design

The experimental design and timeline is outlined in **Fig. 1**. All animals were trained on the rotarod for three days (3 trials per day). Baseline performance was taken on day 0 for body weight, hot-plate and tail-flick assays, and rotarod. Vehicle or ZCZ011 treatment were started on Day 1 and 1 h following treatment animals underwent body weight, hot plate, tail flick, and rotarod tests to assess acute effects of vehicle or ZCZ011. Mice were treated for 14 days with appropriate treatment with continued rotarod assessment (1 trial per day). On day 14, mice undertook another body weight, hot plate, tail flick, and rotarod test to assess chronic effects of vehicle or ZCZ011 treatment. Additionally, mice were habituated to the open field for evaluating locomotor activity. After 24 h (day 15) mice were treated with vehicle or ZCZ011 and 1 h later mice were evaluated in the locomotor activity and elevated plus maze (EPM) tasks. Additionally, the novel object recognition (NOR) task was conducted on Day 16 with either vehicle or ZCZ011 injection administered after NOR training and 1 h prior NOR testing. Lastly, on Day 17, mice received their last treatment and were sacrificed 1 h after. Brains and spinal cord were dissected and snap frozen for further evaluation. The right and left hemisphere of the brain were used for mass spectrometry and western blot analysis, respectively.

### Behavioral procedure

#### Spontaneous heat-evoked nociception

The hot-plate and tail-flick assays were used to evaluate spontaneous heat-evoked nociception with an emphasis on supraspinal- and spinal-related pathways, respectively [124].

For the tail-flick assay, the distal 1/3^rd^ of the tail of each mouse was dipped in a water bath (Thermo Scientific, Precision General-Purpose Water Bath, Model 181, MA, USA) maintained at 56 ± 1°C. A maximum cut-off latency of 10 s was used to prevent tissue damage. The mouse’s latency to flick and remove the tail from the water bath was recorded as a measure of nociception. Following the tail-flick assay, the hot-plate assay was conducted. Each mouse was placed on the hot-plate surface (IITC Inc., MOD 39, CA, USA) within a Plexiglas™ cylinder (15 cm height, 10 cm diameter) to avoid escape. The hot plate was maintained as 55 ± 1°C and a 15 s cut-off latency was used to prevent tissue damage. Mice were removed immediately after withdrawing or licking a paw or jumping and total time was recorded as a measure of nociception.

#### Motor coordination

A rotarod test was used to evaluate motor function and coordination as previously described [125]. The rotarod apparatus (Harvard Apparatus, #76-0770, MA, USA) consists of a raised, rubber-covered rod (30 mm diameter, elevated 18 cm) divided into five sections (50 mm wide each) to allow for simultaneous testing of multiple animals. Mice were trained on the rotarod for three consecutive days (3 trials/day) prior testing. On test days, mice were placed on the rod and allowed to habituate for 1 min before starting the test (1 trial). The accelerating rod is initially rotated at 2 rotations per minute (rpm) and the speed is increased by 1 rpm every 7 s up to 60 rpm which was reached by 7 min. Animals’ performance on the rotarod was continued to be assessed over the 14-day treatment period. The amount of time (s) each animal remained on the rotating rod without falling was recorded for Days 0, 1, and 14.

#### Locomotor activity

Spontaneous motor activity was evaluate using the open field activity chamber (SD Instruments, Photobeam Activity System – Open Field, CA, USA). Mice were habituated to the chamber 24 h prior testing. The activity chamber consists of 16” x 16” cm Plexiglas™ enclosure that is wired with 16 photo-beam cells (each x- and y-axis) connect to a computer console. The photo-beams accurately record the ambulatory movements of the animal contained with the enclosure. On the test day, ambulatory movements (locomotor activity) were recorded for a 5 min period. The habituation and the locomotor activity testing was conducted in a dark room.

#### Elevated plus maze

The elevated plus maze (EPM) task is a common measure of anxiety-like behavior as in mice increased anxiety is related to increased preference for darker and enclosed spaces [126]. Mice were placed in the center of an elevated, plus-shaped maze consisting of two closed-arms with sheltered beige walls (15 cm tall) and two open-arms (unsheltered). All arms were 30 cm long, 5 cm wide, and 38 cm elevated. The EPM was placed under indirect lighting illuminating all four walls consistently between animals. Mice were free to explore the apparatus for 10 min and their behavior was recorded with a video camera (GoPro, Hero 6 Black, v2.10, CA, USA) mounted over the EPM maze. Numerical data were generated from videos by a team of 3 trained experimenters blinded to treatment condition. The time spent in the open arms was used as indices of open space-induced anxiety. Data are presented as percent time spent in the open arms from total exploration time.

#### Novel object recognition task

The novel object recognition (NOR) task is a well-established task to measure object recognition memory which relies upon the natural tendency of mice to explore new objects or stimuli [127, 128]. The NOR task was conducted in the open field activity chamber (SD Instruments, Photobeam Activity System – Open Field, CA, USA) used for the locomotor activity task. The task consisted of three phases: habituation, training, and testing as previously described [128]. The locomotor activity test on Day 15 was used as the habituation phase for NOR training and testing on Day 16. In the training phase, two identical (familiar) objects were presented and mice were left to explore the arena for 10 min. Following the training phase, mice were treated with vehicle or ZCZ011 and returned to the home cage. Testing phase began 60 min after training and treatment. For the testing phase, one of the familiar objects was replaced (randomized for each mouse) by a new (novel) object and mice were allowed to explore the arena for 10 min to assess object recognition memory. Photo-beam breaks were used to record the time that mice spent exploring the familiar and novel object during the session. As mice show an innate preference for novelty, their preference for the novel object was used to quantify successful object recognition memory [127]. The objects used in the task were approximately of equal size but of different shape and color and no natural significance to mice. Total exploration time (s) of both familiar and novel objects were calculated to assess object exploration time. Discrimination between the objects was measured using a discrimination index (DI), calculated as [(time spent exploring the novel object) – (time spent exploring the familiar object)] / total time spent exploring both objects. A discrimination index of 0 indicated no preference, 1 indicates complete preference for the novel object, and –1 indicates complete preference for the familiar object.

### Analysis of endocannabinoids and related lipids

Endogenous cannabinoid ligands, including the two main endocannabinoids *N*-arachidonoylethanolamine (AEA) and 2-arachidonoylglycerol (2-AG), and two minor endocannabinoids, including *N*-oleoylethanolamide (OEA) and *N*-palmitoylethanolamide (PEA), and arachidonic acid (AA), were quantified via ultraperformance liquid chromatography-tandem mass spectrometry (UPLC-MS/MS) from four CNS regions (prefrontal cortex, striatum, cerebellum, and spinal cord) of female and male Tat transgenic mice after completion of behavioral experiments. Following the behavior experiments, mice were returned to the home cage. On the next day, all the animals received the last dose of their respective treatment and sacrificed by rapid decapitation after isoflurane-induced anesthesia after 60 min. The seven CNS regions were dissected from the right brain hemisphere and were snap-frozen in liquid nitrogen. Samples were stored at –80°C until they were processed, and substrates quantified as previously described [129]. Extraction and quantification methods of endocannabinoids and related lipids is as previously described [61].

### Western blot analysis

CNS regions, including prefrontal cortex, striatum, hippocampus, cortex, cerebellum, brain stem from the left hemisphere, and spinal cord samples of female and male Tat transgenic mice were quantified for cannabinoid receptor protein expression, including cannabinoid type 1 and 2 receptors (CB_1_ and CB_2_R), and endocannabinoid degradative enzymes, including fatty acid amide hydrolase (FAAH) and monoacylglycerol lipase (MAGL). Samples were homogenized on ice in an appropriate volume of ice-cold Pierce™ RIPA lysis and extraction buffer (Thermo Scientific, Cat# 89900, USA) with Halt™ phosphatase (Thermo Scientific, Cat# 78420, USA) and protease inhibitor cocktail (Thermo Scientific, Cat# 87786, USA) followed by centrifugation at 10,000 g for 10 min at 4 °C. Protein concentration in the tissue lysates were determined by Pierce™ BCA protein assay kit (Thermo Scientific, Cat# 23227, USA). Protein lysates were suspended in sample buffer containing NuPAGE™ LDS Sample Buffer (Invitrogen™, Cat# NP0007, USA) and XT Reducing Agent (BioRad, Cat# 1610792, USA) in 1:2.5 ratio and denatured at 85°C for 10 min. Equal amounts of protein (20 μg/lane) were resolved in 10% Bis-Tris Criterion™ XT Precast Gels and XT MOPS running buffer (BioRad, Cat# 1610799, USA) at 120 volts for 1.5 h using Criterion™ vertical electrophoresis cell (BioRad, Cat# 1656001, USA). The resolved proteins were transferred from the gel to Immobilon®-P PVDF membranes (Millipore Sigma, Cat# IPVH00010, USA) in 10x Tris/Glycine buffer (BioRad, Cat# 1610734) at 1-4 °C and 100 volts for 1 h using Criterion™ blotter with wire electrodes (BioRad, Cat#1704071, USA). Blots were rinsed with phosphate-buffered saline (PBS), incubated with Intercept® blocking buffer (LI-COR Biosciences, Cat# 727-70001) at room temperature for 1 h followed by overnight incubation with primary antibodies overnight at 4 °C in Intercept® blocking buffer with 0.2% Tween-20. Primary antibodies used in this study were, anti-CB_1_R (rabbit polyclonal; Proteintech, Cat# 17978-1-AP, 1:1000 dilution), anti-CB_1_R (rabbit polyclonal; AbClonal, Cat# A1762, 1:1000 dilution), anti-FAAH (mouse monoclonal; Abcam, Cat# ab54615, 1:1000 dilution), and anti-MAGL (rabbit polyclonal; Abcam, Cat# ab24701, 1:1000 dilution). Anti-GAPDH antibody (mouse monoclonal; Abcam, Cat# ab125247, 1:15,000 dilution) was used as a loading control. Following primary antibody incubation, the blots were washed 3x with PBST (PBS with 0.1% Tween-20) and incubated with IRDye^®^ 680RD Donkey anti-Mouse IgG (LI-COR Biosciences, Cat# 926-68072, 1:15,000 dilution) and IRDye^®^ 800CW Donkey anti-Rabbit IgG (LI-COR Biosciences, Cat# 925-32213, 1:15,000 dilution) secondary antibodies at room temperature for 1 h in Intercept® blocking buffer with 0.2% Tween-20 and 0.01% SDS. The blots were washed 3x with PBST and bands were detected using Odyssey® CLx infrared imaging system (LI-COR Biosciences, USA) and analyzed in Empiria studio® software version 2.3.0 (LI-COR Biosciences, USA). Data presented are normalized to the housekeeping gene GAPDH.

### Statistical analysis

All data are presented as mean ± the standard error of the mean (SEM). Baseline data for behavioral measures (body weight, motor coordination, tail flick, hot plate, and rotarod) were analyzed using a two-way analysis of variances (ANOVAs) with sex (2 levels: females, males) and genotype [2 levels: Tat(–), Tat(+)] as between-subjects factors. Main or interaction effects, were followed by Tukey’s post hoc tests when appropriate. Four-way mixed ANOVAs were conducted on acute and chronic drug treatments with time (2 levels: Day 1, Day 14) as a within-subjects factor, and sex (2 levels: females, males), genotype [2 levels: Tat(–), Tat(+)], and treatment (2 levels: vehicle, ZCZ011) as between-subjects factors. Main or interaction effects for sex were followed up by separate three-way ANOVAs for females and males. Follow-up Tukey’s post hoc tests were conducted when appropriate. Additionally, one-sample *t*-tests were conducted for acute and chronic drug treatments and compared to the baseline (set at 100%). Group comparisons were corrected for multiple comparisons with Tukey’s-adjusted *p* values reported. Behavioral experiments (locomotor activity, elevated plus maze, novel object recognition) and protein quantification data (UPLC-MA/MS and western blot) were analyzed by three-way ANOVAs with sex (2 levels: females, males), genotype [2 levels: Tat(–),Tat(+)], and treatment (2 levels: vehicle, ZCZ011) as between-subjects factors. Main or interaction effects were followed up by separate two-way ANOVAs for females and males. Note, that for rotarod and the novel object recognition task, non-significant interaction trends were also followed up by separate two-way ANOVAs for females and males. Follow-up Tukey’s post hoc tests were conducted when appropriate. An alpha level of *p* ≤ 0.05 was considered significant for all statistical tests. SPSS Statistics 25 (IBM, Chicago, IL) and Prism GraphPad 8.0 (San Diego, CA) was used for data analysis and data graphing, respectively.

## Acknowledgements

The authors would also like to acknowledge the work of animal care technician DeVeda Eubanks for her role in maintaining the welfare of our animals through the studies.

## Supporting Information

**S1_Table. Effect of drug, genotype, and sex on the levels of endocannabinoids and related lipids in nmol/g in four CNS regions.**

**S2_Table. Effect of drug, genotype, and sex on the levels of cannabinoid receptors and degradative enzymes in seven CNS regions.**

**S1_Fig. Original and unedited blots of CB_1_R and CB_2_R expression levels in the prefrontal cortex.** Images show original (**A**) CB_1_R, CB_2_R, and GAPDH and (**B**) FAAH, MAGL, and GAPDH for females and males. Tat(–) vehicle- and ZCZ011-treated mice are represented by lanes 1-4 and 5-8 respectively. Tat(+) vehicle- and ZCZ011-treated mice are represented by lanes 9-12 and 13-16 respectively. M: molecular weights of marker protein (kDa).

**S2_Fig. Original and unedited blots of CB_1_R and CB_2_R expression levels in the striatum.** Images show original (**A**) CB_1_R, CB_2_R, and GAPDH and (**B**) FAAH, MAGL, and GAPDH for females and males. Tat(–) vehicle- and ZCZ011-treated mice are represented by lanes 1-4 and 5-8 respectively. Tat(+) vehicle- and ZCZ011-treated mice are represented by lanes 9-12 and 13-16 respectively. M: molecular weights of marker protein (kDa).

**S3_Fig. Original and unedited blots of CB_1_R and CB_2_R expression levels in the hippocampus.** Images show original (**A**) CB_1_R, CB_2_R, and GAPDH and (**B**) FAAH, MAGL, and GAPDH for females and males. Tat(–) vehicle- and ZCZ011-treated mice are represented by lanes 1-4 and 5-8 respectively. Tat(+) vehicle- and ZCZ011-treated mice are represented by lanes 9-12 and 13-16 respectively. M: molecular weights of marker protein (kDa).

**S4_Fig. Original and unedited blots of CB_1_R and CB_2_R expression levels in the cortex.** Images show original (**A**) CB_1_R, CB_2_R, and GAPDH and (**B**) FAAH, MAGL, and GAPDH for females and males. Tat(–) vehicle- and ZCZ011-treated mice are represented by lanes 1-4 and 5-8 respectively. Tat(+) vehicle- and ZCZ011-treated mice are represented by lanes 9-12 and 13-16 respectively. M: molecular weights of marker protein (kDa).

**S5_Fig. Original and unedited blots of CB_1_R and CB_2_R expression levels in the cerebellum.** Images show original (**A**) CB_1_R, CB_2_R, and GAPDH and (**B**) FAAH, MAGL, and GAPDH for females and males. Tat(–) vehicle- and ZCZ011-treated mice are represented by lanes 1-4 and 5-8 respectively. Tat(+) vehicle- and ZCZ011-treated mice are represented by lanes 9-12 and 13-16 respectively. M: molecular weights of marker protein (kDa). Note: The specks seen in (B) was only seen in the green channel and therefore did not interfere with the quantification of FAAH or GAPDH.

**S6_Fig. Original and unedited blots of CB_1_R and CB_2_R expression levels in the brainstem.** Images show original (**A**) CB_1_R, CB_2_R, and GAPDH and (**B**) FAAH, MAGL, and GAPDH for females and males. Tat(–) vehicle- and ZCZ011-treated mice are represented by lanes 1-4 and 5-8 respectively. Tat(+) vehicle- and ZCZ011-treated mice are represented by lanes 9-12 and respectively. M: molecular weights of marker protein (kDa). Note: The specks seen in (B) was only seen in the green channel and therefore did not interfere with the quantification of FAAH or GAPDH.

**S7_Fig. Original and unedited blots of CB_1_R and CB_2_R expression levels in the spinal cord.** Images show original (**A**) CB_1_R, CB_2_R, and GAPDH and (**B**) FAAH, MAGL, and GAPDH for females and males. Tat(–) vehicle- and ZCZ011-treated mice are represented by lanes 1-4 and 5-8 respectively. Tat(+) vehicle- and ZCZ011-treated mice are represented by lanes 9-12 and 13-16 respectively. M: molecular weights of marker protein (kDa).

## References

1. WHO. Epidemiological fact sheet, HIV statistics, globally and by WHO region 2023 2023 [Available from: https://cdn.who.int/media/docs/default-source/hq-hiv-hepatitis-and-stis-library/j0294-who-hiv-epi-factsheet-v7.pdf

2. Vella S, Schwartlander B, Sow SP, Eholie SP, Murphy RL. The history of antiretroviral therapy and of its implementation in resource-limited areas of the world. AIDS. 2012;26(10):1231–41. doi: 10.1097/QAD.0b013e32835521a3.

3. Harrison KM, Song R, Zhang X. Life expectancy after HIV diagnosis based on national HIV surveillance data from 25 states, United States. J Acquir Immune Defic Syndr. 2010;53(1):124–30. doi: 10.1097/QAI.0b013e3181b563e7.

4. Heaton RK, Franklin DR, Ellis RJ, McCutchan JA, Letendre SL, Leblanc S, et al. HIV-associated neurocognitive disorders before and during the era of combination antiretroviral therapy: differences in rates, nature, and predictors. J Neurovirol. 2011;17(1):3–16. doi: 10.1007/s13365-010-0006-1.

5. Martinez-Picado J, Deeks SG. Persistent HIV-1 replication during antiretroviral therapy. Curr Opin HIV AIDS. 2016;11(4):417–23. doi: 10.1097/COH.0000000000000287.

6. Masliah E, Heaton RK, Marcotte TD, Ellis RJ, Wiley CA, Mallory M, et al. Dendritic injury is a pathological substrate for human immunodeficiency virus-related cognitive disorders. HNRC Group. The HIV Neurobehavioral Research Center. Ann Neurol. 1997;42(6):963–72. doi: 10.1002/ana.410420618.

7. Ellis R, Langford D, Masliah E. HIV and antiretroviral therapy in the brain: neuronal injury and repair. Nat Rev Neurosci. 2007;8(1):33–44. doi: 10.1038/nrn2040.

8. El-Hage N, Rodriguez M, Dever SM, Masvekar RR, Gewirtz DA, Shacka JJ. HIV-1 and morphine regulation of autophagy in microglia: limited interactions in the context of HIV-1 infection and opioid abuse. J Virol. 2015;89(2):1024–35. doi: 10.1128/JVI.02022-14.

9. Sippy BD, Hofman FM, Wallach D, Hinton DR. Increased expression of tumor necrosis factor-alpha receptors in the brains of patients with AIDS. J Acquir Immune Defic Syndr Hum Retrovirol. 1995;10(5):511–21. doi:

10. Wesselingh SL, Takahashi K, Glass JD, McArthur JC, Griffin JW, Griffin DE. Cellular localization of tumor necrosis factor mRNA in neurological tissue from HIV-infected patients by combined reverse transcriptase/polymerase chain reaction in situ hybridization and immunohistochemistry. J Neuroimmunol. 1997;74(1-2):1–8. doi: 10.1016/s0165-5728(96)00160-9.

11. Kovalevich J, Langford D. Neuronal toxicity in HIV CNS disease. Future Virol. 2012;7(7):687–98. doi: 10.2217/fvl.12.57.

12. King JE, Eugenin EA, Buckner CM, Berman JW. HIV tat and neurotoxicity. Microbes Infect. 2006;8(5):1347–57. doi: 10.1016/j.micinf.2005.11.014.

13. Carroll A, Brew B. HIV-associated neurocognitive disorders: recent advances in pathogenesis, biomarkers, and treatment. F1000Res. 2017;6:312. doi: 10.12688/f1000research.10651.1.

14. Rao VR, Ruiz AP, Prasad VR. Viral and cellular factors underlying neuropathogenesis in HIV associated neurocognitive disorders (HAND). AIDS Res Ther. 2014;11:13. doi: 10.1186/1742-6405-11-13.

15. Rappaport J, Joseph J, Croul S, Alexander G, Del Valle L, Amini S, Khalili K. Molecular pathway involved in HIV-1-induced CNS pathology: role of viral regulatory protein, Tat. J Leukoc Biol. 1999;65(4):458–65. doi: 10.1002/jlb.65.4.458.

16. Prendergast MA, Rogers DT, Mulholland PJ, Littleton JM, Wilkins LH, Jr., Self RL, Nath A. Neurotoxic effects of the human immunodeficiency virus type-1 transcription factor Tat require function of a polyamine sensitive-site on the N-methyl-D-aspartate receptor. Brain Res. 2002;954(2):300–7. doi: 10.1016/s0006-8993(02)03360-7.

17. Behnisch T, Francesconi W, Sanna PP. HIV secreted protein Tat prevents long-term potentiation in the hippocampal CA1 region. Brain Res. 2004;1012(1-2):187–9. doi: 10.1016/j.brainres.2004.03.037.

18. Longordo F, Feligioni M, Chiaramonte G, Sbaffi PF, Raiteri M, Pittaluga A. The human immunodeficiency virus-1 protein transactivator of transcription up-regulates N-methyl-D-aspartate receptor function by acting at metabotropic glutamate receptor 1 receptors coexisting on human and rat brain noradrenergic neurones. J Pharmacol Exp Ther. 2006;317(3):1097–105. doi: 10.1124/jpet.105.099630.

19. Zucchini S, Pittaluga A, Brocca-Cofano E, Summa M, Fabris M, De Michele R, et al. Increased excitability in tat-transgenic mice: role of tat in HIV-related neurological disorders. Neurobiol Dis. 2013;55:110–9. doi: 10.1016/j.nbd.2013.02.004.

20. Fitting S, Knapp PE, Zou S, Marks WD, Bowers MS, Akbarali HI, Hauser KF. Interactive HIV-1 Tat and morphine-induced synaptodendritic injury is triggered through focal disruptions in Na(+) influx, mitochondrial instability, and Ca(2)(+) overload. J Neurosci. 2014;34(38):12850–64. doi: 10.1523/JNEUROSCI.5351-13.2014.

21. Bertrand SJ, Mactutus CF, Aksenova MV, Espensen-Sturges TD, Booze RM. Synaptodendritic recovery following HIV Tat exposure: neurorestoration by phytoestrogens. J Neurochem. 2014;128(1):140–51. doi: 10.1111/jnc.12375.

22. Hahn YK, Vo P, Fitting S, Block ML, Hauser KF, Knapp PE. beta-Chemokine production by neural and glial progenitor cells is enhanced by HIV-1 Tat: effects on microglial migration. J Neurochem. 2010;114(1):97–109. doi: 10.1111/j.1471-4159.2010.06744.x.

23. Nath A, Conant K, Chen P, Scott C, Major EO. Transient exposure to HIV-1 Tat protein results in cytokine production in macrophages and astrocytes. A hit and run phenomenon. J Biol Chem. 1999;274(24):17098–102. doi: 10.1074/jbc.274.24.17098.

24. Sheng WS, Hu S, Hegg CC, Thayer SA, Peterson PK. Activation of human microglial cells by HIV-1 gp41 and Tat proteins. Clin Immunol. 2000;96(3):243–51. doi: 10.1006/clim.2000.4905.

25. Zou S, Fitting S, Hahn YK, Welch SP, El-Hage N, Hauser KF, Knapp PE. Morphine potentiates neurodegenerative effects of HIV-1 Tat through actions at mu-opioid receptor-expressing glia. Brain. 2011;134(Pt 12):3616–31. doi: 10.1093/brain/awr281.

26. Gannon P, Khan MZ, Kolson DL. Current understanding of HIV-associated neurocognitive disorders pathogenesis. Curr Opin Neurol. 2011;24(3):275–83. doi: 10.1097/WCO.0b013e32834695fb.

27. Matsuda LA, Lolait SJ, Brownstein MJ, Young AC, Bonner TI. Structure of a cannabinoid receptor and functional expression of the cloned cDNA. Nature. 1990;346(6284):561–4. doi: 10.1038/346561a0.

28. Howlett AC, Barth F, Bonner TI, Cabral G, Casellas P, Devane WA, et al. International Union of Pharmacology. XXVII. Classification of cannabinoid receptors. Pharmacol Rev. 2002;54(2):161–202. doi: 10.1124/pr.54.2.161.

29. Navarrete M, Araque A. Endocannabinoids potentiate synaptic transmission through stimulation of astrocytes. Neuron. 2010;68(1):113–26. doi: 10.1016/j.neuron.2010.08.043.

30. Oliveira da Cruz JF, Robin LM, Drago F, Marsicano G, Metna-Laurent M. Astroglial type-1 cannabinoid receptor (CB1): A new player in the tripartite synapse. Neuroscience. 2016;323:35–42. doi: 10.1016/j.neuroscience.2015.05.002.

31. Harkany T, Mackie K, Doherty P. Wiring and firing neuronal networks: endocannabinoids take center stage. Curr Opin Neurobiol. 2008;18(3):338–45. doi: 10.1016/j.conb.2008.08.007.

32. Chevaleyre V, Takahashi KA, Castillo PE. Endocannabinoid-mediated synaptic plasticity in the CNS. Annu Rev Neurosci. 2006;29:37–76. doi: 10.1146/annurev.neuro.29.051605.112834.

33. Chiarlone A, Bellocchio L, Blazquez C, Resel E, Soria-Gomez E, Cannich A, et al. A restricted population of CB1 cannabinoid receptors with neuroprotective activity. Proc Natl Acad Sci U S A. 2014;111(22):8257–62. doi: 10.1073/pnas.1400988111.

34. Marsicano G, Goodenough S, Monory K, Hermann H, Eder M, Cannich A, et al. CB1 cannabinoid receptors and on-demand defense against excitotoxicity. Science. 2003;302(5642):84–8. doi: 10.1126/science.1088208.

35. Rossi S, Furlan R, De Chiara V, Muzio L, Musella A, Motta C, et al. Cannabinoid CB1 receptors regulate neuronal TNF-alpha effects in experimental autoimmune encephalomyelitis. Brain Behav Immun. 2011;25(6):1242–8. doi: 10.1016/j.bbi.2011.03.017.

36. Derkinderen P, Valjent E, Toutant M, Corvol JC, Enslen H, Ledent C, et al. Regulation of extracellular signal-regulated kinase by cannabinoids in hippocampus. J Neurosci. 2003;23(6):2371–82. doi: 10.1523/JNEUROSCI.23-06-02371.2003.

37. Hampson RE, Miller F, Palchik G, Deadwyler SA. Cannabinoid receptor activation modifies NMDA receptor mediated release of intracellular calcium: implications for endocannabinoid control of hippocampal neural plasticity. Neuropharmacology. 2011;60(6):944–52. doi: 10.1016/j.neuropharm.2011.01.039.

38. Li Q, Yan H, Wilson WA, Swartzwelder HS. Modulation of NMDA and AMPA-mediated synaptic transmission by CB1 receptors in frontal cortical pyramidal cells. Brain Res. 2010;1342:127–37. doi: 10.1016/j.brainres.2010.04.029.

39. Liu Q, Bhat M, Bowen WD, Cheng J. Signaling pathways from cannabinoid receptor-1 activation to inhibition of N-methyl-D-aspartic acid mediated calcium influx and neurotoxicity in dorsal root ganglion neurons. J Pharmacol Exp Ther. 2009;331(3):1062–70. doi: 10.1124/jpet.109.156216.

40. Sanchez-Blazquez P, Rodriguez-Munoz M, Garzon J. The cannabinoid receptor 1 associates with NMDA receptors to produce glutamatergic hypofunction: implications in psychosis and schizophrenia. Front Pharmacol. 2014;4:169. doi: 10.3389/fphar.2013.00169.

41. Pertwee RG. Elevating endocannabinoid levels: pharmacological strategies and potential therapeutic applications. Proc Nutr Soc. 2014;73(1):96–105. doi: 10.1017/S0029665113003649.

42. Scotter EL, Abood ME, Glass M. The endocannabinoid system as a target for the treatment of neurodegenerative disease. Br J Pharmacol. 2010;160(3):480–98. doi: 10.1111/j.1476-5381.2010.00735.x.

43. Cosenza-Nashat MA, Bauman A, Zhao ML, Morgello S, Suh HS, Lee SC. Cannabinoid receptor expression in HIV encephalitis and HIV-associated neuropathologic comorbidities. Neuropathol Appl Neurobiol. 2011;37(5):464–83. doi: 10.1111/j.1365-2990.2011.01177.x.

44. Benito C, Kim WK, Chavarria I, Hillard CJ, Mackie K, Tolon RM, et al. A glial endogenous cannabinoid system is upregulated in the brains of macaques with simian immunodeficiency virus-induced encephalitis. Journal of Neuroscience. 2005;25(10):2530–6. doi: Doi 10.1523/Jneurosci.3923-04.2005.

45. Pryce G, Baker D. Potential control of multiple sclerosis by cannabis and the endocannabinoid system. CNS Neurol Disord Drug Targets. 2012;11(5):624–41. doi: 10.2174/187152712801661310.

46. Fagan SG, Campbell VA. The influence of cannabinoids on generic traits of neurodegeneration. Br J Pharmacol. 2014;171(6):1347–60. doi: 10.1111/bph.12492.

47. Hutcheson DM, Tzavara ET, Smadja C, Valjent E, Roques BP, Hanoune J, Maldonado R. Behavioural and biochemical evidence for signs of abstinence in mice chronically treated with delta-9-tetrahydrocannabinol. Br J Pharmacol. 1998;125(7):1567–77. doi: 10.1038/sj.bjp.0702228.

48. Justinova Z, Tanda G, Redhi GH, Goldberg SR. Self-administration of delta9-tetrahydrocannabinol (THC) by drug naive squirrel monkeys. Psychopharmacology (Berl). 2003;169(2):135–40. doi: 10.1007/s00213-003-1484-0.

49. Cooper ZD, Haney M. Actions of delta-9-tetrahydrocannabinol in cannabis: relation to use, abuse, dependence. Int Rev Psychiatry. 2009;21(2):104–12. doi: 10.1080/09540260902782752.

50. Kenakin T. Analytical pharmacology and allosterism: the importance of quantifying drug parameters in drug discovery. Drug Discov Today Technol. 2013;10(2):e229–35. doi: 10.1016/j.ddtec.2012.07.006.

51. Kenakin T, Strachan RT. PAM-Antagonists: A Better Way to Block Pathological Receptor Signaling? Trends Pharmacol Sci. 2018;39(8):748–65. doi: 10.1016/j.tips.2018.05.001.

52. Price MR, Baillie GL, Thomas A, Stevenson LA, Easson M, Goodwin R, et al. Allosteric modulation of the cannabinoid CB1 receptor. Mol Pharmacol. 2005;68(5):1484–95. doi: 10.1124/mol.105.016162.

53. Ross RA. Allosterism and cannabinoid CB(1) receptors: the shape of things to come. Trends Pharmacol Sci. 2007;28(11):567–72. doi: 10.1016/j.tips.2007.10.006.

54. Pertwee RG. The therapeutic potential of drugs that target cannabinoid receptors or modulate the tissue levels or actions of endocannabinoids. AAPS J. 2005;7(3):E625–54. doi: 10.1208/aapsj070364.

55. Ignatowska-Jankowska BM, Baillie GL, Kinsey S, Crowe M, Ghosh S, Owens RA, et al. A Cannabinoid CB1 Receptor-Positive Allosteric Modulator Reduces Neuropathic Pain in the Mouse with No Psychoactive Effects. Neuropsychopharmacology. 2015;40(13):2948–59. doi: 10.1038/npp.2015.148.

56. Slivicki RA, Iyer V, Mali SS, Garai S, Thakur GA, Crystal JD, Hohmann AG. Positive Allosteric Modulation of CB(1) Cannabinoid Receptor Signaling Enhances Morphine Antinociception and Attenuates Morphine Tolerance Without Enhancing Morphine-Induced Dependence or Reward. Front Mol Neurosci. 2020;13:54. doi: 10.3389/fnmol.2020.00054.

57. Green HM, Finlay DB, Ross RA, Greig IR, Duffull SB, Glass M. In Vitro Characterization of 6-Methyl-3-(2-nitro-1-(thiophen-2-yl)ethyl)-2-phenyl-1H-indole (ZCZ011) at the Type 1 Cannabinoid Receptor: Allosteric Agonist or Allosteric Modulator? ACS Pharmacol Transl Sci. 2022;5(12):1279–91. doi: 10.1021/acsptsci.2c00160.

58. Cairns EA, Szczesniak AM, Straiker AJ, Kulkarni PM, Pertwee RG, Thakur GA, et al. The In Vivo Effects of the CB(1)-Positive Allosteric Modulator GAT229 on Intraocular Pressure in Ocular Normotensive and Hypertensive Mice. J Ocul Pharmacol Ther. 2017;33(8):582–90. doi: 10.1089/jop.2017.0037.

59. Datta U, Kelley LK, Middleton JW, Gilpin NW. Positive allosteric modulation of the cannabinoid type-1 receptor (CB1R) in periaqueductal gray (PAG) antagonizes anti-nociceptive and cellular effects of a mu-opioid receptor agonist in morphine-withdrawn rats. Psychopharmacology (Berl). 2020;237(12):3729–39. doi: 10.1007/s00213-020-05650-5.

60. Trexler KR, Eckard ML, Kinsey SG. CB(1) positive allosteric modulation attenuates Delta(9)-THC withdrawal and NSAID-induced gastric inflammation. Pharmacol Biochem Behav. 2019;177:27–33. doi: 10.1016/j.pbb.2018.12.009.

61. Yadav-Samudrala BJ, Gorman BL, Dodson H, Ramineni S, Wallace ED, Peace MR, et al. Effects of acute Delta(9)-tetrahydrocannabinol on behavior and the endocannabinoid system in HIV-1 Tat transgenic female and male mice. Brain Res. 2024;1822:148638. doi: 10.1016/j.brainres.2023.148638.

62. Dodu JC, Moncayo RK, Damaj MI, Schlosburg JE, Akbarali HI, O’Brien LD, et al. The Cannabinoid Receptor Type 1 Positive Allosteric Modulator ZCZ011 Attenuates Naloxone-Precipitated Diarrhea and Weight Loss in Oxycodone-Dependent Mice. J Pharmacol Exp Ther. 2022;380(1):1–14. doi: 10.1124/jpet.121.000723.

63. Laprairie RB, Bagher AM, Rourke JL, Zrein A, Cairns EA, Kelly MEM, et al. Positive allosteric modulation of the type 1 cannabinoid receptor reduces the signs and symptoms of Huntington’s disease in the R6/2 mouse model. Neuropharmacology. 2019;151:1–12. doi: 10.1016/j.neuropharm.2019.03.033.

64. Hernandez AR, Truckenbrod LM, Campos KT, Williams SA, Burke SN. Sex differences in age-related impairments vary across cognitive and physical assessments in rats. Behav Neurosci. 2020;134(2):69–81. doi: 10.1037/bne0000352.

65. Slivicki RA, Xu Z, Kulkarni PM, Pertwee RG, Mackie K, Thakur GA, Hohmann AG. Positive Allosteric Modulation of Cannabinoid Receptor Type 1 Suppresses Pathological Pain Without Producing Tolerance or Dependence. Biol Psychiatry. 2018;84(10):722–33. doi: 10.1016/j.biopsych.2017.06.032.

66. Deuis JR, Dvorakova LS, Vetter I. Methods Used to Evaluate Pain Behaviors in Rodents. Front Mol Neurosci. 2017;10:284. doi: 10.3389/fnmol.2017.00284.

67. Bardo MT, Hughes RA. Exposure to a nonfunctional hot plate as a factor in the assessment of morphine-induced analgesia and analgesic tolerance in rats. Pharmacol Biochem Behav. 1979;10(4):481–5. doi: 10.1016/0091-3057(79)90221-1.

68. Gamble GD, Milne RJ. Repeated exposure to sham testing procedures reduces reflex withdrawal and hot-plate latencies: attenuation of tonic descending inhibition? Neurosci Lett. 1989;96(3):312–7. doi: 10.1016/0304-3940(89)90397-2.

69. Lai YY, Chan SH. Shortened pain response time following repeated algesiometric tests in rats. Physiol Behav. 1982;28(6):1111–3. doi: 10.1016/0031-9384(82)90184-6.

70. Gunn A, Bobeck EN, Weber C, Morgan MM. The influence of non-nociceptive factors on hot-plate latency in rats. J Pain. 2011;12(2):222–7. doi: 10.1016/j.jpain.2010.06.011.

71. Deng L, Guindon J, Cornett BL, Makriyannis A, Mackie K, Hohmann AG. Chronic cannabinoid receptor 2 activation reverses paclitaxel neuropathy without tolerance or cannabinoid receptor 1-dependent withdrawal. Biol Psychiatry. 2015;77(5):475–87. doi: 10.1016/j.biopsych.2014.04.009.

72. Kinsey SG, Wise LE, Ramesh D, Abdullah R, Selley DE, Cravatt BF, Lichtman AH. Repeated low-dose administration of the monoacylglycerol lipase inhibitor JZL184 retains cannabinoid receptor type 1-mediated antinociceptive and gastroprotective effects. J Pharmacol Exp Ther. 2013;345(3):492–501. doi: 10.1124/jpet.112.201426.

73. Schlosburg JE, Blankman JL, Long JZ, Nomura DK, Pan B, Kinsey SG, et al. Chronic monoacylglycerol lipase blockade causes functional antagonism of the endocannabinoid system. Nat Neurosci. 2010;13(9):1113–9. doi: 10.1038/nn.2616.

74. Roebuck AJ, Greba Q, Smolyakova AM, Alaverdashvili M, Marks WN, Garai S, et al. Positive allosteric modulation of type 1 cannabinoid receptors reduces spike-and-wave discharges in Genetic Absence Epilepsy Rats from Strasbourg. Neuropharmacology. 2021;190:108553. doi: 10.1016/j.neuropharm.2021.108553.

75. McElroy DL, Roebuck AJ, Scott GA, Greba Q, Garai S, Denovan-Wright EM, et al. Antipsychotic potential of the type 1 cannabinoid receptor positive allosteric modulator GAT211: preclinical in vitro and in vivo studies. Psychopharmacology (Berl). 2021;238(4):1087–98. doi: 10.1007/s00213-020-05755-x.

76. Carlsson M, Carlsson A. The NMDA antagonist MK-801 causes marked locomotor stimulation in monoamine-depleted mice. J Neural Transm. 1989;75(3):221–6. doi: 10.1007/BF01258633.

77. Moghaddam B, Krystal JH. Capturing the angel in "angel dust": twenty years of translational neuroscience studies of NMDA receptor antagonists in animals and humans. Schizophr Bull. 2012;38(5):942–9. doi: 10.1093/schbul/sbs075.

78. Zhao X, Fan Y, Vann PH, Wong JM, Sumien N, He JJ. Long-term HIV-1 Tat Expression in the Brain Led to Neurobehavioral, Pathological, and Epigenetic Changes Reminiscent of Accelerated Aging. Aging Dis. 2020;11(1):93–107. doi: 10.14336/AD.2019.0323.

79. Yadav-Samudrala BJ, Gorman BL, Barmada KM, Ravula HP, Huguely CJ, Wallace ED, et al. Effects of acute cannabidiol on behavior and the endocannabinoid system in HIV-1 Tat transgenic female and male mice. Frontiers in Neuroscience. 2024;18. doi: 10.3389/fnins.2024.1358555.

80. Hahn YK, Paris JJ, Lichtman AH, Hauser KF, Sim-Selley LJ, Selley DE, Knapp PE. Central HIV-1 Tat exposure elevates anxiety and fear conditioned responses of male mice concurrent with altered mu-opioid receptor-mediated G-protein activation and beta-arrestin 2 activity in the forebrain. Neurobiol Dis. 2016;92(Pt B):124–36. doi: 10.1016/j.nbd.2016.01.014.

81. Hermes DJ, Jacobs IR, Key MC, League AF, Yadav-Samudrala BJ, Xu C, et al. Escalating morphine dosing in HIV-1 Tat transgenic mice with sustained Tat exposure reveals an allostatic shift in neuroinflammatory regulation accompanied by increased neuroprotective non-endocannabinoid lipid signaling molecules and amino acids. J Neuroinflammation. 2020;17(1):345. doi: 10.1186/s12974-020-01971-6.

82. Liu X, Wang H, Zhu Z, Zhang L, Cao J, Zhang L, et al. Exploring bridge symptoms in HIV-positive people with comorbid depressive and anxiety disorders. Bmc Psychiatry. 2022;22(1):448. doi: 10.1186/s12888-022-04088-7.

83. Mannes ZL, Dunne EM, Ferguson EG, Cook RL, Ennis N. Symptoms of generalized anxiety disorder as a risk factor for substance use among adults living with HIV. Aids Care. 2021;33(5):623–32. doi: 10.1080/09540121.2020.1808163.

84. Whetten K, Reif S, Whetten R, Murphy-McMillan LK. Trauma, mental health, distrust, and stigma among HIV-positive persons: implications for effective care. Psychosom Med. 2008;70(5):531–8. doi: 10.1097/PSY.0b013e31817749dc.

85. Joshi CR, Stacy S, Sumien N, Ghorpade A, Borgmann K. Astrocyte HIV-1 Tat Differentially Modulates Behavior and Brain MMP/TIMP Balance During Short and Prolonged Induction in Transgenic Mice. Front Neurol. 2020;11:593188. doi: 10.3389/fneur.2020.593188.

86. Paris JJ, Fenwick J, McLaughlin JP. Progesterone protects normative anxiety-like responding among ovariectomized female mice that conditionally express the HIV-1 regulatory protein, Tat, in the CNS. Horm Behav. 2014;65(5):445–53. doi: 10.1016/j.yhbeh.2014.04.001.

87. Salahuddin MF, Mahdi F, Paris JJ. HIV-1 Tat Dysregulates the Hypothalamic-Pituitary-Adrenal Stress Axis and Potentiates Oxycodone-Mediated Psychomotor and Anxiety-Like Behavior of Male Mice. Int J Mol Sci. 2020;21(21). doi: 10.3390/ijms21218212.

88. Qrareya AN, Mahdi F, Kaufman MJ, Ashpole NM, Paris JJ. HIV-1 Tat promotes age-related cognitive, anxiety-like, and antinociceptive impairments in female mice that are moderated by aging and endocrine status. Geroscience. 2021;43(1):309–27. doi: 10.1007/s11357-020-00268-z.

89. Franklin DR, Bowker KR, Blumenthal TD. Anxiety and prepulse inhibition of acoustic startle in a normative sample: The importance of signal-to-noise ratio. Personality and Individual Differences. 2009;46:369–73. doi:

90. Cysique LA, Maruff P, Brew BJ. Prevalence and pattern of neuropsychological impairment in human immunodeficiency virus-infected/acquired immunodeficiency syndrome (HIV/AIDS) patients across pre- and post-highly active antiretroviral therapy eras: a combined study of two cohorts. J Neurovirol. 2004;10(6):350–7. doi: 10.1080/13550280490521078.

91. Garvey LJ, Yerrakalva D, Winston A. Correlations between computerized battery testing and a memory questionnaire for identification of neurocognitive impairment in HIV type 1-infected subjects on stable antiretroviral therapy. AIDS Res Hum Retroviruses. 2009;25(8):765–9. doi: 10.1089/aid.2008.0292.

92. Marks WD, Paris JJ, Schier CJ, Denton MD, Fitting S, McQuiston AR, et al. HIV-1 Tat causes cognitive deficits and selective loss of parvalbumin, somatostatin, and neuronal nitric oxide synthase expressing hippocampal CA1 interneuron subpopulations. J Neurovirol. 2016;22(6):747–62. doi: 10.1007/s13365-016-0447-2.

93. Carey AN, Sypek EI, Singh HD, Kaufman MJ, McLaughlin JP. Expression of HIV-Tat protein is associated with learning and memory deficits in the mouse. Behav Brain Res. 2012;229(1):48–56. doi: 10.1016/j.bbr.2011.12.019.

94. Cyrenne DL, Brown GR. Ontogeny of sex differences in response to novel objects from adolescence to adulthood in lister-hooded rats. Dev Psychobiol. 2011;53(7):670–6. doi: 10.1002/dev.20542.

95. Frick KM, Gresack JE. Sex differences in the behavioral response to spatial and object novelty in adult C57BL/6 mice. Behav Neurosci. 2003;117(6):1283–91. doi: 10.1037/0735-7044.117.6.1283.

96. Swinton MK, Sundermann EE, Pedersen L, Nguyen JD, Grelotti DJ, Taffe MA, et al. Alterations in Brain Cannabinoid Receptor Levels Are Associated with HIV-Associated Neurocognitive Disorders in the ART Era: Implications for Therapeutic Strategies Targeting the Endocannabinoid System. Viruses. 2021;13(9). doi: 10.3390/v13091742.

97. Hermes DJ, Yadav-Samudrala BJ, Xu C, Paniccia JE, Meeker RB, Armstrong ML, et al. GPR18 drives FAAH inhibition-induced neuroprotection against HIV-1 Tat-induced neurodegeneration. Exp Neurol. 2021;341:113699. doi: 10.1016/j.expneurol.2021.113699.

98. Hermes DJ, Xu C, Poklis JL, Niphakis MJ, Cravatt BF, Mackie K, et al. Neuroprotective effects of fatty acid amide hydrolase catabolic enzyme inhibition in a HIV-1 Tat model of neuroAIDS. Neuropharmacology. 2018;141:55–65. doi: 10.1016/j.neuropharm.2018.08.013.

99. Xu C, Yadav-Samudrala BJ, Xu C, Nath B, Mistry T, Jiang W, et al. Inhibitory Neurotransmission Is Sex-Dependently Affected by Tat Expression in Transgenic Mice and Suppressed by the Fatty Acid Amide Hydrolase Enzyme Inhibitor PF3845 via Cannabinoid Type-1 Receptor Mechanisms. Cells. 2022;11(5). doi: 10.3390/cells11050857.

100. Jacobs IR, Xu C, Hermes DJ, League AF, Xu C, Nath B, et al. Inhibitory Control Deficits Associated with Upregulation of CB(1)R in the HIV-1 Tat Transgenic Mouse Model of Hand. J Neuroimmune Pharmacol. 2019;14(4):661–78. doi: 10.1007/s11481-019-09867-w.

101. Yadav-Samudrala BJ, Ravula HP, Barmada KM, Dodson H, Poklis JL, Ignatowska-Jankowska BM, et al. Acute Effects of Monoacylglycerol Lipase Inhibitor ABX1431 on Neuronal Hyperexcitability, Nociception, Locomotion, and the Endocannabinoid System in HIV-1 Tat Male Mice. Cannabis Cannabinoid Res. 2024. doi: 10.1089/can.2023.0247.

102. Duarte EAC, Benevides ML, Martins ALP, Duarte EP, Weller ABS, de Azevedo LOC, et al. Female sex is strongly associated with cognitive impairment in HIV infection. Neurol Sci. 2021;42(5):1853–60. doi: 10.1007/s10072-020-04705-x.

103. Rubin LH, Neigh GN, Sundermann EE, Xu Y, Scully EP, Maki PM. Sex Differences in Neurocognitive Function in Adults with HIV: Patterns, Predictors, and Mechanisms. Curr Psychiatry Rep. 2019;21(10):94. doi: 10.1007/s11920-019-1089-x.

104. Maki PM, Martin-Thormeyer E. HIV, cognition and women. Neuropsychol Rev. 2009;19(2):204–14. doi: 10.1007/s11065-009-9093-2.

105. Sundermann EE, Heaton RK, Pasipanodya E, Moore RC, Paolillo EW, Rubin LH, et al. Sex differences in HIV-associated cognitive impairment. AIDS. 2018;32(18):2719–26. doi: 10.1097/QAD.0000000000002012.

106. Santinelli L, Ceccarelli G, Borrazzo C, Innocenti GP, Frasca F, Cavallari EN, et al. Sex-related differences in markers of immune activation in virologically suppressed HIV-infected patients. Biol Sex Differ. 2020;11(1):23. doi: 10.1186/s13293-020-00302-x.

107. Ziegler S, Altfeld M. Sex differences in HIV-1-mediated immunopathology. Curr Opin HIV AIDS. 2016;11(2):209–15. doi: 10.1097/COH.0000000000000237.

108. Yadav-Samudrala BJ, Gorman BL, Barmada K, Ravula HP, Huguely CJ, Wallace ED, et al. Effects of acute cannabidiol on behavior and the endocannabinoid system in HIV-1 Tat transgenic female and male mice. Frontiers in Neuroscience. 2024;18. doi: 10.3389/fnins.2024.1358555.

109. Craft RM, Marusich JA, Wiley JL. Sex differences in cannabinoid pharmacology: a reflection of differences in the endocannabinoid system? Life Sci. 2013;92(8-9):476–81. doi: 10.1016/j.lfs.2012.06.009.

110. Romero J, Garcia-Palomero E, Berrendero F, Garcia-Gil L, Hernandez ML, Ramos JA, Fernandez-Ruiz JJ. Atypical location of cannabinoid receptors in white matter areas during rat brain development. Synapse. 1997;26(3):317–23. doi: 10.1002/(SICI)1098-2396(199707)26:3<317::AID-SYN12>3.0.CO;2-S.

111. Xing G, Carlton J, Jiang X, Wen J, Jia M, Li H. Differential Expression of Brain Cannabinoid Receptors between Repeatedly Stressed Males and Females may Play a Role in Age and Gender-Related Difference in Traumatic Brain Injury: Implications from Animal Studies. Front Neurol. 2014;5:161. doi: 10.3389/fneur.2014.00161.

112. Lopez HH. Cannabinoid-hormone interactions in the regulation of motivational processes. Horm Behav. 2010;58(1):100–10. doi: 10.1016/j.yhbeh.2009.10.005.

113. Gonzalez S, Bisogno T, Wenger T, Manzanares J, Milone A, Berrendero F, et al. Sex steroid influence on cannabinoid CB(1) receptor mRNA and endocannabinoid levels in the anterior pituitary gland. Biochem Biophys Res Commun. 2000;270(1):260–6. doi: 10.1006/bbrc.2000.2406.

114. Rodriguez de Fonseca F, Cebeira M, Ramos JA, Martin M, Fernandez-Ruiz JJ. Cannabinoid receptors in rat brain areas: sexual differences, fluctuations during estrous cycle and changes after gonadectomy and sex steroid replacement. Life Sci. 1994;54(3):159–70. doi: 10.1016/0024-3205(94)00585-0.

115. Maccarrone M. Metabolism of the Endocannabinoid Anandamide: Open Questions after 25 Years. Front Mol Neurosci. 2017;10:166. doi: 10.3389/fnmol.2017.00166.

116. Xing G, Carlton J, Zhang L, Jiang X, Fullerton C, Li H, Ursano R. Cannabinoid receptor expression and phosphorylation are differentially regulated between male and female cerebellum and brain stem after repeated stress: implication for PTSD and drug abuse. Neurosci Lett. 2011;502(1):5–9. doi: 10.1016/j.neulet.2011.05.013.

117. Ong WY, Mackie K. A light and electron microscopic study of the CB1 cannabinoid receptor in primate brain. Neuroscience. 1999;92(4):1177–91. doi: 10.1016/s0306-4522(99)00025-1.

118. Fuerte-Hortigon A, Goncalves J, Zeballos L, Masa R, Gomez-Nieto R, Lopez DE. Distribution of the Cannabinoid Receptor Type 1 in the Brain of the Genetically Audiogenic Seizure-Prone Hamster GASH/Sal. Front Behav Neurosci. 2021;15:613798. doi: 10.3389/fnbeh.2021.613798.

119. Suarez J, Bermudez-Silva FJ, Mackie K, Ledent C, Zimmer A, Cravatt BF, de Fonseca FR. Immunohistochemical description of the endogenous cannabinoid system in the rat cerebellum and functionally related nuclei. J Comp Neurol. 2008;509(4):400–21. doi: 10.1002/cne.21774.

120. Santa-Cecilia FV, Leite CA, Del-Bel E, Raisman-Vozari R. The Neuroprotective Effect of Doxycycline on Neurodegenerative Diseases. Neurotox Res. 2019;35(4):981–6. doi: 10.1007/s12640-019-00015-z.

121. Bruce-Keller AJ, Turchan-Cholewo J, Smart EJ, Geurin T, Chauhan A, Reid R, et al. Morphine causes rapid increases in glial activation and neuronal injury in the striatum of inducible HIV-1 Tat transgenic mice. Glia. 2008;56(13):1414–27. doi: 10.1002/glia.20708.

122. Chauhan A, Turchan J, Pocernich C, Bruce-Keller A, Roth S, Butterfield DA, et al. Intracellular human immunodeficiency virus Tat expression in astrocytes promotes astrocyte survival but induces potent neurotoxicity at distant sites via axonal transport. J Biol Chem. 2003;278(15):13512–9. doi: 10.1074/jbc.M209381200.

123. Eckard ML, Trexler KR, Kotson BT, Anderson KG, Kinsey SG. Precipitated Delta9-THC withdrawal reduces motivation for sucrose reinforcement in mice. Pharmacol Biochem Behav. 2020;195:172966. doi: 10.1016/j.pbb.2020.172966.

124. Singh P, Kongara K, Harding D, Ward N, Dukkipati VSR, Johnson C, Chambers P. Comparison of electroencephalographic changes in response to acute electrical and thermal stimuli with the tail flick and hot plate test in rats administered with opiorphin. BMC Neurol. 2018;18(1):43. doi: 10.1186/s12883-018-1047-y.

125. Jones BJ, Roberts DJ. The quantiative measurement of motor inco-ordination in naive mice using an acelerating rotarod. J Pharm Pharmacol. 1968;20(4):302–4. doi: 10.1111/j.2042-7158.1968.tb09743.x.

126. Shoji H, Miyakawa T. Effects of test experience, closed-arm wall color, and illumination level on behavior and plasma corticosterone response in an elevated plus maze in male C57BL/6J mice: a challenge against conventional interpretation of the test. Mol Brain. 2021;14(1):34. doi: 10.1186/s13041-020-00721-2.

127. Ennaceur A. One-trial object recognition in rats and mice: methodological and theoretical issues. Behav Brain Res. 2010;215(2):244–54. doi: 10.1016/j.bbr.2009.12.036.

128. Lueptow LM. Novel Object Recognition Test for the Investigation of Learning and Memory in Mice. J Vis Exp. 2017(126). doi: 10.3791/55718.

129. League AF, Gorman BL, Hermes DJ, Johnson CT, Jacobs IR, Yadav-Samudrala BJ, et al. Monoacylglycerol Lipase Inhibitor MJN110 Reduces Neuronal Hyperexcitability, Restores Dendritic Arborization Complexity, and Regulates Reward-Related Behavior in Presence of HIV-1 Tat. Front Neurol. 2021;12:651272. doi: 10.3389/fneur.2021.651272.

