## Supplemental Information for "Cannabinoid receptor 1 positive allosteric modulator ZCZ011 shows differential effects on behavior and the endocannabinoid system in HIV-1 Tat transgenic female and male mice"

**S1_Table. Effect of drug, genotype, and sex on the levels of endocannabinoids and related lipids in nmol/g in four CNS regions [a].**

| CNS Region | Lipids nm/mg | Sex | Genotype | Vehicle mean ± SEM | ZCZ011 mean ± SEM | Genotype Effect *p* | Sex Effect *p* | Drug Effect *p* | Genotype x Drug *p* | Genotype x Sex *p* | Sex x Drug p |
|---|---|---|---|---|---|---|---|---|---|---|---|
| PFC | AEA | Female | Tat (+) | 0.06 ± 0.02 | 0.12 ± 0.04 | 0.84 | 0.89 | 0.29 | 0.12 | 0.27 | 0.30 |
| | | | Tat (–) | 0.10 ± 0.01 | 0.12 ± 0.04 | | | | | | |
| | | Male | Tat (+) | 0.09 ± 0.01 | 0.14 ± 0.04 | | | | | | |
| | | | Tat (–) | 0.10 ± 0.03 | 0.06 ± 0.01 | | | | | | |
| | 2-AG | Female | Tat (+) | 3.56 ± 0.48 | 3.21 ± 1.00 | 0.34 | 0.84 | 0.41 | 0.10 | 0.95 | 0.18 |
| | | | Tat (–) | 1.44 ± 0.10 | 4.28 ± 1.57 | | | | | | |
| | | Male | Tat (+) | 3.63 ± 0.78 | 2.99 ± 0.65 | | | | | | |
| | | | Tat (–) | 2.70 ± 0.55 | 2.73 ± 0.48 | | | | | | |
| | PEA | Female | Tat (+) | 0.96 ± 0.24 | 1.67 ± 0.53 | 0.86 | 0.83 | 0.35 | 0.43 | 0.47 | 0.07 |
| | | | Tat (–) | 1.06 ± 0.12 | 1.88 ± 0.75 | | | | | | |
| | | Male | Tat (+) | 1.34 ± 0.23 | 1.58 ± 0.30 | | | | | | |
| | | | Tat (–) | 1.58 ± 0.35 | 0.85 ± 0.08 | | | | | | |
| | OEA | Female | Tat (+) | 0.68 ± 0.22 | 1.17 ± 0.33 | 0.65 | 0.99 | 0.32 | 0.37 | 0.25 | 0.11 |
| | | | Tat (–) | 0.80 ± 0.05 | 1.33 ± 0.51 | | | | | | |
| | | Male | Tat (+) | 1.03 ± 1.18 | 1.27 ± 0.25 | | | | | | |
| | | | Tat (–) | 1.09 ± 0.26 | 0.60 ± 0.06 | | | | | | |
| | AA | Female | Tat (+) | 401.18 ± 138.88 | 630.27 ± 196.22 | 0.23 | 0.81 | 0.27 | 0.22 | 0.29 | 0.25 |
| | | | Tat (–) | 422.21 ± 24.24 | 585.58 ± 138.63 | | | | | | |
| | | Male | Tat (+) | 498.14 ± 76.64 | 676.38 ± 172.73 | | | | | | |
| | | | Tat (–) | 484.27 ± 128.88 | 299.19 ± 35.70 | | | | | | |
| Str | AEA | Female | Tat (+) | 0.06 ± 0.02 | 0.05 ± 0.003 | 0.53 | **0.004** | 0.23 | 0.53 | 0.16 | 0.89 |
| | | | Tat (–) | 0.07 ± 0.01 | 0.06 ± 0.08 | | | | | | |
| | | Male | Tat (+) | 0.09 ± 0.10 | 0.09 ± 0.01 | | | | | | |
| | | | Tat (–) | 0.08 ± 0.01 | 0.07 ± 0.005 | | | | | | |
| | 2-AG | Female | Tat (+) | 11.86 ± 5.32 | 14.16 ± 7.18 | 0.42 | 0.10 | 0.87 | 0.53 | 0.49 | 0.70 |
| | | | Tat (–) | 8.07 ± 5.17 | 8.99 ± 3.36 | | | | | | |
| | | Male | Tat (+) | 4.94 ± 0.68 | 6.95 ± 2.70 | | | | | | |
| | | | Tat (–) | 7.32 ± 3.91 | 3.92 ± 0.41 | | | | | | |
| | PEA | Female | Tat (+) | 1.60 ± 0.27 | 1.70 ± 0.27 | 0.30 | **<0.001** | 0.46 | 0.48 | 0.07 | 0.75 |
| | | | Tat (–) | 2.18 ± 0.40 | 1.55 ± 0.34 | | | | | | |
| | | Male | Tat (+) | 3.21 ± 0.25 | 3.11 ± 0.68 | | | | | | |
| | | | Tat (–) | 2.46 ± 0.27 | 2.36 ± 0.13 | | | | | | |
| | OEA | Female | Tat (+) | 1.28 ± 0.23 | 1.48 ± 0.25 | 0.32 | **0.005** | 0.62 | 0.46 | 0.08 | 0.80 |
| | | | Tat (–) | 1.78 ± 0.24 | 1.30 ± 0.33 | | | | | | |
| | | Male | Tat (+) | 2.39 ± 0.29 | 2.30 ± 0.46 | | | | | | |

| | | | | | | | | | | | |
|---|---|---|---|---|---|---|---|---|---|---|---|
| | | | Tat (–) | 1.79 ± 0.12 | 1.79 ± 0.15 | | | | | | |
| | AA | Female | Tat (+) | 501.07 ± 133.43 | 410.10 ± 22.41 | 0.55 | **0.04** | 0.12 | 0.86 | 0.43 | 0.69 |
| | | | Tat (–) | 533.64 ± 86.14 | 399.03 ± 56.91 | | | | | | |
| | | Male | Tat (+) | 655.84 ± 46.32 | 583.86 ± 115.32 | | | | | | |
| | | | Tat (–) | 572.43 ± 66.42 | 508.45 ± 49.71 | | | | | | |
| Crb | AEA | Female | Tat (+) | 0.04 ± 0.01 | 0.04 ± 0.003 | 0.35 | 0.32 | **0.02** | **0.01** | 0.06 | 0.89 |
| | | | Tat (–) | 0.04 ± 0.002 | 0.07 ± 0.02 | | | | | | |
| | | Male | Tat (+) | 0.04 ± 0.004 | 0.05 ± 0.004 | | | | | | |
| | | | Tat (–) | 0.03 ± 0.004 | 0.05 ± 0.004 | | | | | | |
| | 2-AG | Female | Tat (+) | 11.29 ± 1.95 | 6.96 ± 0.85 | 0.29 | 0.24 | 0.29 | **0.03** | **0.01** | 0.43 |
| | | | Tat (–) | 9.67 ± 1.09 | 12.18 ± 1.49 | | | | | | |
| | | Male | Tat (+) | 17.73 ± 3.97 | 12.01 ± 1.80 | | | | | | |
| | | | Tat (–) | 9.27 ± 3.08 | 9.34 ± 0.92 | | | | | | |
| | PEA | Female | Tat (+) | 0.95 ± 0.21 | 0.66 ± 0.85 | 0.84 | 0.14 | 0.21 | **0.03** | 0.35 | 0.88 |
| | | | Tat (–) | 0.66 ± 0.13 | 1.24 ± 0.30 | | | | | | |
| | | Male | Tat (+) | 0.71 ± 0.26 | 0.75 ± 0.10 | | | | | | |
| | | | Tat (–) | 0.47 ± 0.15 | 0.79 ± 0.11 | | | | | | |
| | OEA | Female | Tat (+) | 1.29 ± 0.16 | 1.08 ± 0.09 | 0.57 | 0.33 | **0.03** | **0.009** | 0.11 | 0.94 |
| | | | Tat (–) | 1.08 ± 0.11 | 1.83 ± 0.36 | | | | | | |
| | | Male | Tat (+) | 1.23 ± 0.19 | 1.29 ± 0.08 | | | | | | |
| | | | Tat (–) | 0.87 ± 0.14 | 1.39 ± 0.10 | | | | | | |
| | AA | Female | Tat (+) | 706.06 ± 77.57 | 585.90 ± 65.28 | 0.09 | **0.01** | 0.29 | **0.01** | **0.02** | 0.72 |
| | | | Tat (–) | 731.06 ± 63.77 | 998.91 ± 107.82 | | | | | | |
| | | Male | Tat (+) | 654.94 ± 82.28 | 604.69 ± 44.53 | | | | | | |
| | | | Tat (–) | 530.02 ± 68.15 | 655.24 ± 61.46 | | | | | | |
| SC | AEA | Female | Tat (+) | 0.02 ± 0.004 | 0.03 ± 0.006 | 0.40 | 0.80 | 0.74 | **0.05** | 0.37 | 0.26 |
| | | | Tat (–) | 0.03 ± 0.003 | 0.02 ± 0.004 | | | | | | |
| | | Male | Tat (+) | 0.02 ± 0.002 | 0.03 ± 0.003 | | | | | | |
| | | | Tat (–) | 0.04 ± 0.02 | 0.01 ± 0.003 | | | | | | |
| | 2-AG | Female | Tat (+) | 25.81 ± 5.66 | 26.34 ± 5.34 | 0.75 | 0.26 | 0.74 | 0.40 | 0.35 | 0.47 |
| | | | Tat (–) | 19.73 ± 1.83 | 22.35 ± 2.28 | | | | | | |
| | | Male | Tat (+) | 25.09 ± 6.73 | 28.64 ± 2.36 | | | | | | |
| | | | Tat (–) | 35.36 ± 10.18 | 23.36 ± 5.46 | | | | | | |
| | PEA | Female | Tat (+) | 2.50 ± 0.34 | 3.05 ± 0.72 | 0.62 | 0.78 | 0.65 | 0.09 | 0.10 | 0.25 |
| | | | Tat (–) | 2.22 ± 0.33 | 2.28 ± 0.44 | | | | | | |
| | | Male | Tat (+) | 1.65 ± 0.24 | 2.20 ± 0.23 | | | | | | |
| | | | Tat (–) | 3.84 ± 1.37 | 1.90 ± 0.17 | | | | | | |
| | OEA | Female | Tat (+) | 1.58 ± 0.18 | 2.21 ± 9.46 | 0.54 | 0.73 | 0.88 | **0.05** | 0.12 | 0.21 |
| | | | Tat (–) | 1.64 ± 0.22 | 1.62 ± 0.28 | | | | | | |
| | | Male | Tat (+) | 1.18 ± 0.17 | 1.56 ± 0.15 | | | | | | |
| | | | Tat (–) | 2.54 ± 9.85 | 1.39 ± 0.89 | | | | | | |

| | | | | | | | | | | | |
|---|---|---|---|---|---|---|---|---|---|---|---|
| | AA | Female | Tat (+) | 450.31 ± 51.83 | 596.88 ± 111.80 | 0.33 | **0.01** | 0.67 | 0.20 | 0.07 | 0.22 |
| | | | Tat (–) | 509.50 ± 57.85 | 447.97 ± 69.76 | | | | | | |
| | | Male | Tat (+) | 324.37 ± 37.38 | 269.74 ± 46.42 | | | | | | |
| | | | Tat (–) | 504.92 ± 117.44 | 384.74 ± 50.25 | | | | | | |

[a]Levels of N-arachidonoylethanolamine (AEA), 2-arachidonoylglycerol (2-AG), palmitoylethanolamide (PEA), oleoylethanolamide (OEA), arachidonic acid (AA) in the prefrontal cortex, striatum, cerebellum, and spinal cord of Tat(–) and Tat(+) female and male mice exposed to chronic 10 mg/kg ZCZ011 or vehicle expressed as mean ± SEM. A three-way ANOVE for each lipid molecule was conducted with drug, genotype, and sex as between-subjects factors. Red bolded values denote significant differences at $p$ <0.05; $N$ = 32(16F).

**S2_Table. Effect of drug, genotype, and sex on the levels of cannabinoid receptors and degradative enzymes in seven CNS regions [a].**

| CNS Region | Receptors and enzymes | Sex | Genotype | Vehicle mean ± SEM | ZCZ011 mean ± SEM | Genotype Effect *p* | Sex Effect *p* | Drug Effect *p* | Genotype x Drug *p* | Genotype x Sex *p* | Sex x Drug *p* |
|---|---|---|---|---|---|---|---|---|---|---|---|
| PFC | $CB_1R$ | Female | Tat (+) | 0.40 ± 0.13 | 0.49 ± 0.08 | 0.17 | 0.85 | 0.25 | 0.16 | 0.07 | 0.17 |
| | | | Tat (–) | 0.54 ± 0.01 | 0.42 ± 0.04 | | | | | | |
| | | Male | Tat (+) | 0.44 ± 0.13 | 0.76 ± 0.20 | | | | | | |
| | | | Tat (–) | 0.31 ± 0.04 | 0.39 ± 0.04 | | | | | | |
| | $CB_2R$ | Female | Tat (+) | 1.11 ± 0.12 | 0.65 ± 0.12 | **0.04** | **0.02** | 0.31 | 0.40 | 0.14 | 0.10 |
| | | | Tat (–) | 0.94 ± 0.16 | 0.86 ± 0.13 | | | | | | |
| | | Male | Tat (+) | 1.35 ± 0.39 | 2.22 ± 0.55 | | | | | | |
| | | | Tat (–) | 1.00 ± 0.17 | 1.16 ± 0.14 | | | | | | |
| | FAAH | Female | Tat (+) | 0.74 ± 0.10 | 0.54 ± 0.12 | 0.27 | **0.03** | 0.06 | 0.26 | **0.01** | 0.09 |
| | | | Tat (–) | 2.46 ± 0.76 | 0.78 ± 0.04 | | | | | | |
| | | Male | Tat (+) | 2.01 ± 0.42 | 1.81 ± 0.31 | | | | | | |
| | | | Tat (–) | 1.44 ± 0.26 | 1.56 ± 0.31 | | | | | | |
| | MAGL | Female | Tat (+) | 0.32 ± 0.01 | 0.33 ± 0.06 | 0.06 | **<0.001** | **0.05** | **0.04** | **0.05** | **0.04** |
| | | | Tat (–) | 1.15 ± 0.40 | 0.30 ± 0.02 | | | | | | |
| | | Male | Tat (+) | 0.03 ± 0.01 | 0.03 ± 0.004 | | | | | | |
| | | | Tat (–) | 0.18 ± 0.006 | 0.02 ± 0.006 | | | | | | |
| Str | $CB_1R$ | Female | Tat (+) | 0.34 ± 0.04 | 0.50 ± 0.07 | 0.12 | **<0.001** | 0.96 | **0.003** | 0.45 | 0.61 |
| | | | Tat (–) | 0.45 ± 0.02 | 0.28 ± 0.02 | | | | | | |
| | | Male | Tat (+) | 0.11 ± 0.01 | 0.11 ± 0.01 | | | | | | |
| | | | Tat (–) | 0.07 ± 0.11 | 0.11 ± 0.02 | | | | | | |
| | $CB_2R$ | Female | Tat (+) | 0.97 ± 0.02 | 1.27 ± 0.12 | 0.46 | **<0.001** | 0.61 | 0.08 | 0.38 | 0.47 |
| | | | Tat (–) | 1.28 ± 0.06 | 0.96 ± 0.09 | | | | | | |
| | | Male | Tat (+) | 0.74 ± 0.04 | 0.74 ± 0.08 | | | | | | |
| | | | Tat (–) | 0.55 ± 0.07 | 0.72 ± 0.14 | | | | | | |
| | FAAH | Female | Tat (+) | 0.20 ± 0.02 | 0.18 ± 0.02 | 0.93 | **<0.001** | 0.91 | **<0.001** | **0.03** | 0.11 |
| | | | Tat (–) | 0.25 ± 0.01 | 0.22 ± 0.04 | | | | | | |
| | | Male | Tat (+) | 0.24 ± 0.01 | 0.48 ± 0.04 | | | | | | |
| | | | Tat (–) | 0.40 ± 0.004 | 0.24 ± 0.03 | | | | | | |
| | MAGL | Female | Tat (+) | 0.09 ± 0.01 | 0.09 ± 0.01 | 0.78 | **<0.001** | 0.46 | **0.03** | 0.15 | 0.64 |
| | | | Tat (–) | 0.06 ± 0.006 | 0.07 ± 0.01 | | | | | | |
| | | Male | Tat (+) | 0.13 ± 0.01 | 0.21 ± 0.02 | | | | | | |
| | | | Tat (–) | 0.20 ± 0.03 | 0.16 ± 0.03 | | | | | | |

| | | | | | | | | | | | |
|---|---|---|---|---|---|---|---|---|---|---|---|
| Hip | $CB_1R$ | Female | Tat (+) | 0.19 ± 0.01 | 0.13 ± 0.003 | 0.14 | **<0.001** | **<0.001** | **0.04** | **0.002** | **0.04** |
| | | | Tat (–) | 0.21 ± 0.03 | 0.18 ± 0.03 | | | | | | |
| | | Male | Tat (+) | 0.47 ± 0.04 | 0.29 ± 0.04 | | | | | | |
| | | | Tat (–) | 0.29 ± 0.01 | 0.27 ± 0.03 | | | | | | |
| | $CB_2R$ | Female | Tat (+) | 0.99 ± 0.05 | 0.82 ± 0.15 | 0.91 | **<0.001** | **0.008** | 0.53 | **0.01** | 0.78 |
| | | | Tat (–) | 1.23 ± 0.13 | 0.96 ± 0.09 | | | | | | |
| | | Male | Tat (+) | 0.87 ± 0.07 | 0.54 ± 0.04 | | | | | | |
| | | | Tat (–) | 0.52 ± 0.04 | 0.48 ± 0.08 | | | | | | |
| | FAAH | Female | Tat (+) | 0.65 ± 0.16 | 0.64 ± 0.06 | 0.53 | 0.39 | 0.13 | 0.60 | 0.30 | 0.79 |
| | | | Tat (–) | 1.14 ± 0.21 | 0.22 ± 0.04 | | | | | | |
| | | Male | Tat (+) | 1.14 ± 0.24 | 0.80 ± 0.21 | | | | | | |
| | | | Tat (–) | 1.03 ± 0.47 | 0.76 ± 0.19 | | | | | | |
| | MAGL | Female | Tat (+) | 0.59 ± 0.13 | 0.71 ± 0.04 | 0.63 | 0.95 | 0.68 | 0.81 | 0.74 | 0.40 |
| | | | Tat (–) | 0.66 ± 0.12 | 0.07 ± 0.01 | | | | | | |
| | | Male | Tat (+) | 0.80 ± 0.13 | 0.55 ± 0.10 | | | | | | |
| | | | Tat (–) | 0.58 ± 0.24 | 0.61 ± 0.10 | | | | | | |
| Ctx | $CB_1R$ | Female | Tat (+) | 0.07 ± 0.01 | 0.07 ± 0.006 | **0.01** | **<0.001** | 0.21 | 0.42 | **0.002** | **0.006** |
| | | | Tat (–) | 0.09 ± 0.02 | 0.06 ± 0.01 | | | | | | |
| | | Male | Tat (+) | 0.23 ± 0.01 | 0.26 ± 0.02 | | | | | | |
| | | | Tat (–) | 0.17 ± 0.01 | 0.21 ± 0.01 | | | | | | |
| | $CB_2R$ | Female | Tat (+) | 1.71 ± 0.34 | 1.15 ± 0.11 | **0.01** | **<0.001** | 0.23 | 0.34 | **0.02** | 0.29 |
| | | | Tat (–) | 2.2 ± 0.36 | 2.12 ± 0.26 | | | | | | |
| | | Male | Tat (+) | 0.63 ± 0.03 | 0.56 ± 0.06 | | | | | | |
| | | | Tat (–) | 0.59 ± 0.06 | 0.64 ± 0.05 | | | | | | |
| | FAAH | Female | Tat (+) | 2.58 ± 0.70 | 0.89 ± 0.14 | 0.97 | **0.008** | 0.64 | 0.18 | 0.93 | 0.77 |
| | | | Tat (–) | 1.20 ± 0.33 | 2.21 ± 1.60 | | | | | | |
| | | Male | Tat (+) | 0.37 ± 0.04 | 0.43 ± 0.02 | | | | | | |
| | | | Tat (–) | 0.56 ± 0.09 | 0.35 ± 0.04 | | | | | | |
| | MAGL | Female | Tat (+) | 2.12 ± 0.57 | 0.81 ± 0.12 | 0.73 | **<0.001** | 0.19 | 0.16 | 0.69 | 0.19 |
| | | | Tat (–) | 1.24 ± 0.28 | 1.36 ± 0.62 | | | | | | |
| | | Male | Tat (+) | 0.15 ± 0.01 | 0.23 ± 0.04 | | | | | | |
| | | | Tat (–) | 0.24 ± 0.01 | 0.16 ± 0.02 | | | | | | |
| Crb | $CB_1R$ | Female | Tat (+) | 0.27 ± 0.01 | 0.27 ± 0.01 | 0.55 | **<0.001** | 0.62 | 0.57 | **0.02** | 0.43 |
| | | | Tat (–) | 0.36 ± 0.02 | 0.29 ± 0.03 | | | | | | |
| | | Male | Tat (+) | 0.14 ± 0.03 | 0.18 ± 0.07 | | | | | | |
| | | | Tat (–) | 0.08 ± 0.01 | 0.16 ± 0.01 | | | | | | |
| | $CB_2R$ | Female | Tat (+) | 0.90 ± 0.03 | 0.90 ± 0.06 | 0.34 | 0.98 | 0.35 | 0.61 | 0.21 | 0.18 |
| | | | Tat (–) | 1.18 ± 0.09 | 1.09 ± 0.04 | | | | | | |
| | | Male | Tat (+) | 0.99 ± 0.20 | 1.08 ± 0.34 | | | | | | |
| | | | Tat (–) | 0.79 ± 0.10 | 1.20 ± 0.04 | | | | | | |
| | FAAH | Female | Tat (+) | 0.29 ± 0.02 | 0.24 ± 0.02 | **0.01** | **<0.001** | 0.26 | **0.03** | 0.56 | 0.65 |

| | | | | | | | | | | | |
|---|---|---|---|---|---|---|---|---|---|---|---|
| | | | Tat (–) | 0.49 ± 0.02 | 0.33 ± 0.04 | | | | | | |
| | | Male | Tat (+) | 0.54 ± 0.07 | 0.74 ± 0.09 | | | | | | |
| | | | Tat (–) | 1.01 ± 0.17 | 0.72 ± 0.17 | | | | | | |
| | MAGL | Female | Tat (+) | 0.06 ± 0.002 | 0.06 ± 0.002 | **0.01** | **<0.001** | 0.71 | 0.08 | 0.08 | 0.70 |
| | | | Tat (–) | 0.07 ± 0.005 | 0.07 ± 0.003 | | | | | | |
| | | Male | Tat (+) | 0.14 ± 0.02 | 0.20 ± 0.03 | | | | | | |
| | | | Tat (–) | 0.29 ± 0.04 | 0.21 ± 0.06 | | | | | | |
| BS | $CB_1R$ | Female | Tat (+) | 0.39 ± 0.03 | 0.39 ± 0.09 | 0.57 | 0.43 | 0.47 | **0.05** | 0.40 | 0.91 |
| | | | Tat (–) | 0.13 ± 0.03 | 0.29 ± 0.02 | | | | | | |
| | | Male | Tat (+) | 0.54 ± 0.21 | 0.22 ± 0.06 | | | | | | |
| | | | Tat (–) | 0.14 ± 0.03 | 0.68 ± 0.43 | | | | | | |
| | $CB_2R$ | Female | Tat (+) | 1.11 ± 0.05 | 1.17 ± 0.33 | 0.669 | 0.32 | 0.70 | 0.08 | 0.48 | 0.84 |
| | | | Tat (–) | 0.65 ± 0.02 | 0.74 ± 0.02 | | | | | | |
| | | Male | Tat (+) | 1.82 ± 0.64 | 0.69 ± 0.19 | | | | | | |
| | | | Tat (–) | 0.56 ± 0.11 | 2.15 ± 1.35 | | | | | | |
| | FAAH | Female | Tat (+) | 0.07 ± 0.02 | 0.33 ± 0.08 | 0.29 | **0.01** | 0.54 | 0.06 | **0.05** | 0.25 |
| | | | Tat (–) | 0.28 ± 0.03 | 0.02 ± 0.01 | | | | | | |
| | | Male | Tat (+) | 0.25 ± 0.03 | 0.19 ± 0.06 | | | | | | |
| | | | Tat (–) | 0.24 ± 0.04 | 0.23 ± 0.08 | | | | | | |
| | MAGL | Female | Tat (+) | 0.30 ± 0.02 | 0.36 ± 0.03 | **0.001** | **<0.001** | **0.01** | 0.11 | 0.54 | 0.52 |
| | | | Tat (–) | 0.07 ± 0.005 | 0.25 ± 0.04 | | | | | | |
| | | Male | Tat (+) | 0.22 ± 0.03 | 0.29 ± 0.02 | | | | | | |
| | | | Tat (–) | 0.14 ± 0.02 | 0.23 ± 0.04 | | | | | | |
| SC | $CB_1R$ | Female | Tat (+) | 0.39 ± 0.09 | 0.47 ± 0.04 | 0.94 | **<0.001** | 0.55 | 0.40 | **<0.001** | 0.25 |
| | | | Tat (–) | 0.24 ± 0.02 | 0.31 ± 0.01 | | | | | | |
| | | Male | Tat (+) | 0.09 ± 0.02 | 0.13 ± 0.02 | | | | | | |
| | | | Tat (–) | 0.31 ± 0.10 | 0.23 ± 0.04 | | | | | | |
| | $CB_2R$ | Female | Tat (+) | 0.68 ± 0.09 | 0.63 ± 0.04 | 0.27 | **0.006** | 0.94 | 0.58 | **0.002** | 0.94 |
| | | | Tat (–) | 0.34 ± 0.04 | 0.42 ± 0.02 | | | | | | |
| | | Male | Tat (+) | 0.50 ± 0.06 | 0.69 ± 0.14 | | | | | | |
| | | | Tat (–) | 1.21 ± 0.35 | 1.02 ± 0.20 | | | | | | |
| | FAAH | Female | Tat (+) | 0.10 ± 0.07 | 0.18 ± 0.04 | **0.006** | **<0.001** | 0.66 | 0.24 | 0.11 | 0.64 |
| | | | Tat (–) | 0.28 ± 0.06 | 0.19 ± 0.02 | | | | | | |
| | | Male | Tat (+) | 0.34 ± 0.03 | 0.47 ± 0.16 | | | | | | |
| | | | Tat (–) | 0.72 ± 0.12 | 0.71 ± 0.15 | | | | | | |
| | MAGL | Female | Tat (+) | 0.31 ± 0.07 | 0.36 ± 0.03 | **0.03** | **<0.001** | 0.21 | 0.28 | **<0.001** | 0.21 |
| | | | Tat (–) | 0.19 ± 0.04 | 0.14 ± 0.03 | | | | | | |
| | | Male | Tat (+) | 0.35 ± 0.12 | 0.34 ± 0.10 | | | | | | |
| | | | Tat (–) | 0.68 ± 0.10 | 1.20 ± 0.32 | | | | | | |

[a]Levels of cannabinoid type 1 and 2 receptors ($CB_1R$ and $CB_2R$) and degradative enzymes fatty acid amide hydrolase (FAAH) and monoacylglycerol (MAGL in the prefrontal cortex, striatum, hippocampus, cortex, cerebellum, brainstem, and spinal cord of Tat(–) and Tat(+) female and male mice exposed to chronic 10 mg/kg ZCZ011 or vehicle expressed as mean ± SEM. A three-way ANOVE for each protein was conducted with drug, genotype, and sex as between-subjects factors. Red bolded values denote significant differences at $p < 0.05$; $N = 32$(16F).

**Western blot – Raw images**

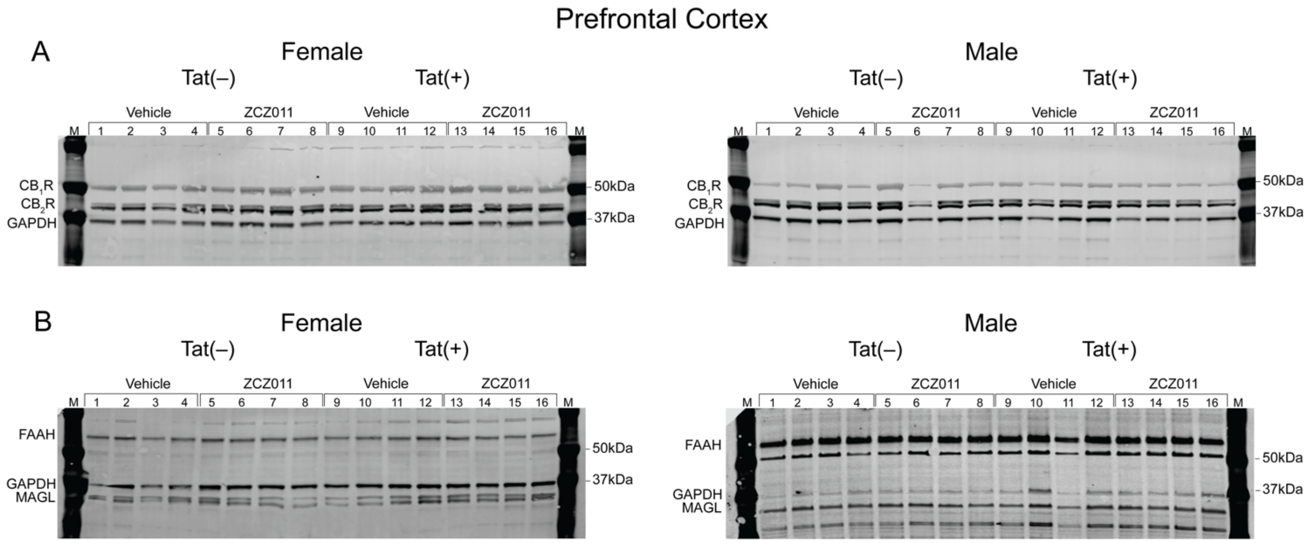

**S1_Fig. Original and unedited blots of $CB_1R$ and $CB_2R$ expression levels in the prefrontal cortex.** Images show original (**A**) $CB_1R$, $CB_2R$, and GAPDH and (**B**) FAAH, MAGL, and GAPDH for females and males. Tat(–) vehicle- and ZCZ011-treated mice are represented by lanes 1-4 and 5-8 respectively. Tat(+) vehicle- and ZCZ011-treated mice are represented by lanes 9-12 and 13-16 respectively. M: molecular weights of marker protein (kDa).

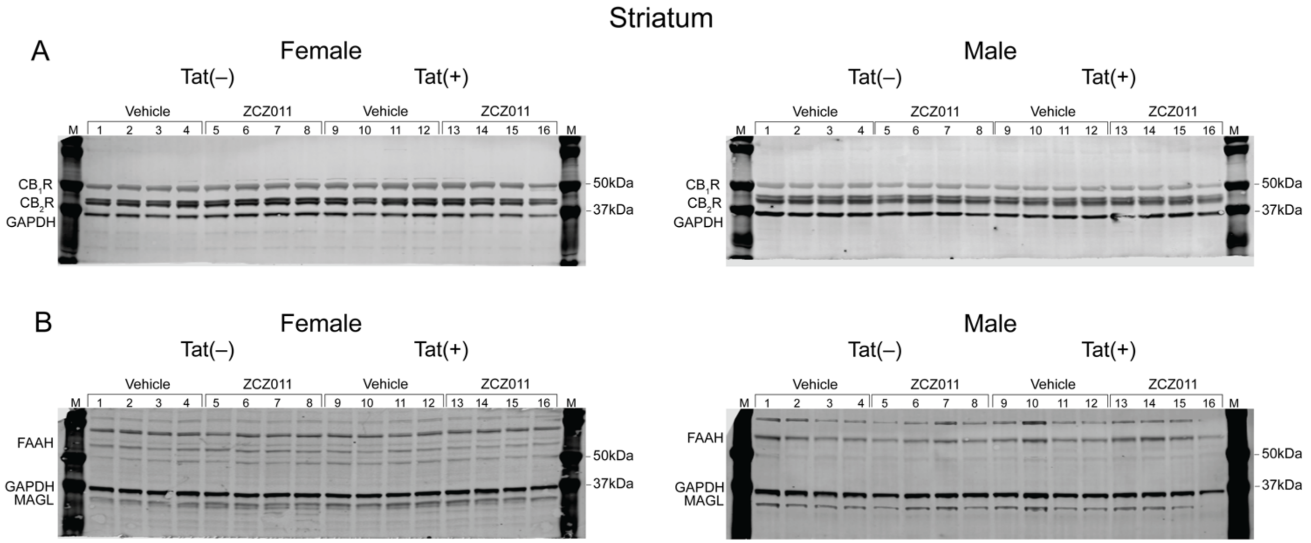

**S2_Fig. Original and unedited blots of $CB_1R$ and $CB_2R$ expression levels in the striatum.** Images show original (**A**) $CB_1R$, $CB_2R$, and GAPDH and (**B**) FAAH, MAGL, and GAPDH for females and males. Tat(–) vehicle- and ZCZ011-treated mice are represented by lanes 1-4 and 5-8 respectively. Tat(+) vehicle- and ZCZ011-treated mice are represented by lanes 9-12 and 13-16 respectively. M: molecular weights of marker protein (kDa).

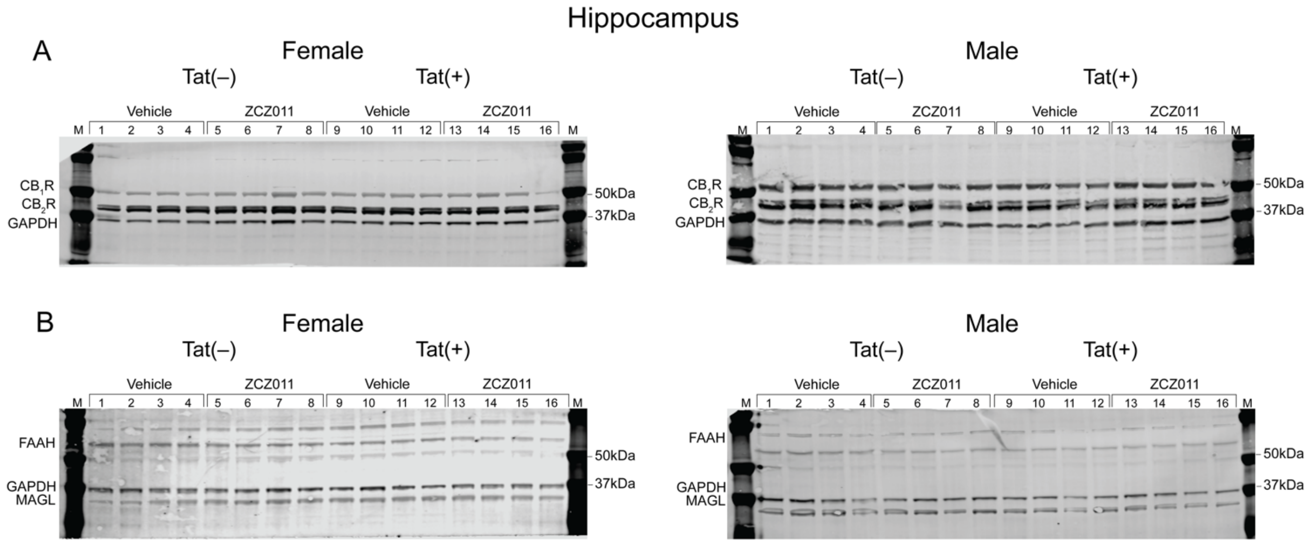

**S3_Fig. Original and unedited blots of $CB_1R$ and $CB_2R$ expression levels in the hippocampus.** Images show original (**A**) $CB_1R$, $CB_2R$, and GAPDH and (**B**) FAAH, MAGL, and GAPDH for females and males. Tat(–) vehicle- and ZCZ011-treated mice are represented by lanes 1-4 and 5-8 respectively. Tat(+) vehicle- and ZCZ011-treated mice are represented by lanes 9-12 and 13-16 respectively. M: molecular weights of marker protein (kDa).

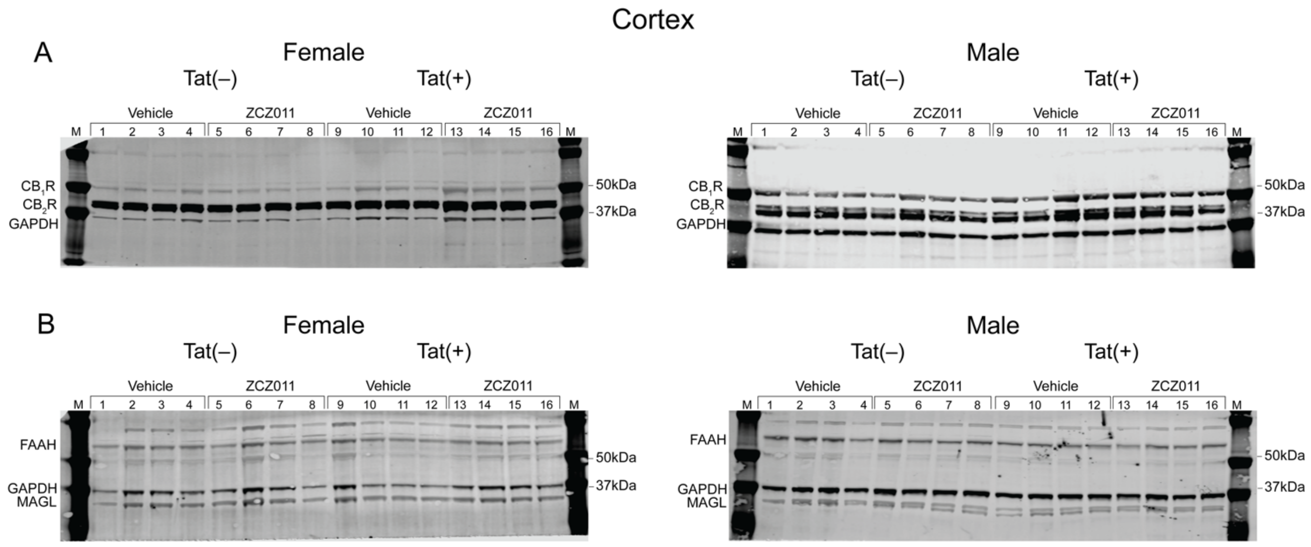

**S4_Fig. Original and unedited blots of $CB_1R$ and $CB_2R$ expression levels in the cortex.** Images show original (**A**) $CB_1R$, $CB_2R$, and GAPDH and (**B**) FAAH, MAGL, and GAPDH for females and males. Tat(–) vehicle- and ZCZ011-treated mice are represented by lanes 1-4 and 5-8 respectively. Tat(+) vehicle- and ZCZ011-treated mice are represented by lanes 9-12 and 13-16 respectively. M: molecular weights of marker protein (kDa).

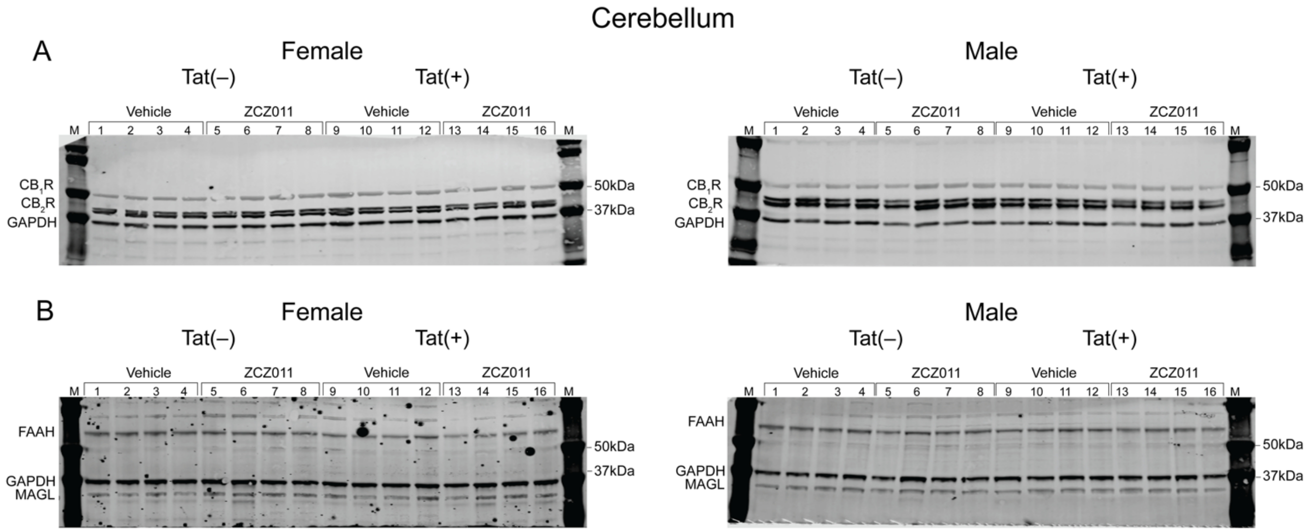

**S5_Fig. Original and unedited blots of $CB_1R$ and $CB_2R$ expression levels in the cerebellum.** Images show original (**A**) $CB_1R$, $CB_2R$, and GAPDH and (**B**) FAAH, MAGL, and GAPDH for females and males. Tat(–) vehicle- and ZCZ011-treated mice are represented by lanes 1-4 and 5-8 respectively. Tat(+) vehicle- and ZCZ011-treated mice are represented by lanes 9-12 and 13-16 respectively. M: molecular weights of marker protein (kDa). Note: The specks seen in (B) was only seen in the green channel and therefore did not interfere with the quantification of FAAH or GAPDH.

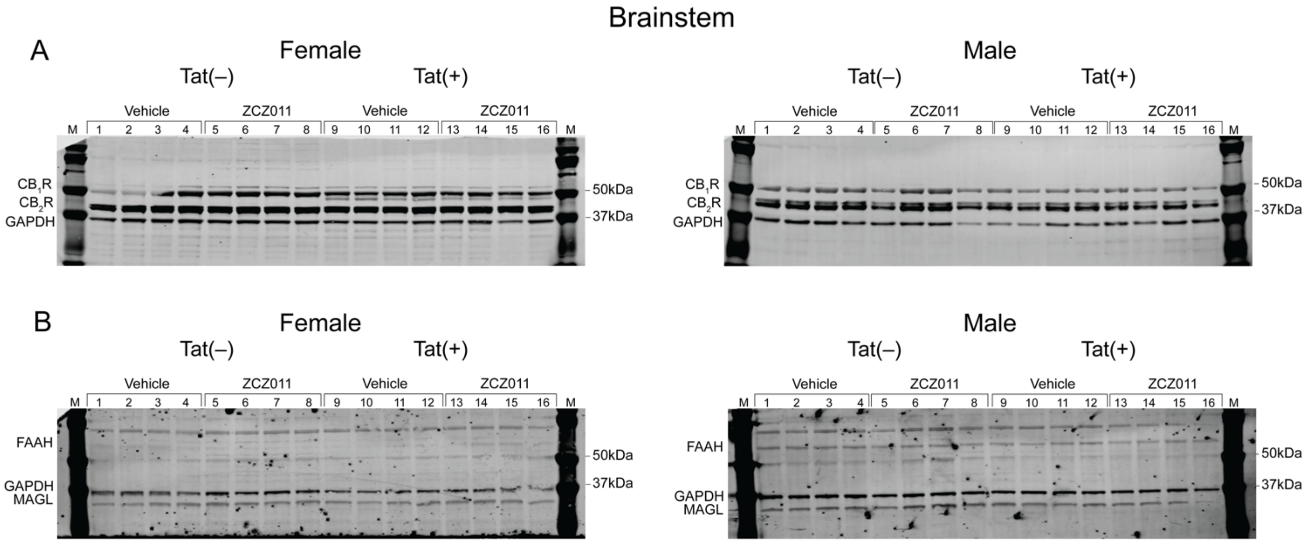

**S6_Fig. Original and unedited blots of $CB_1R$ and $CB_2R$ expression levels in the brainstem.** Images show original (**A**) $CB_1R$, $CB_2R$, and GAPDH and (**B**) FAAH, MAGL, and GAPDH for females and males. Tat(–) vehicle- and ZCZ011-treated mice are represented by lanes 1-4 and 5-8 respectively. Tat(+) vehicle- and ZCZ011-treated mice are represented by lanes 9-12 and 13-16 respectively. M: molecular weights of marker protein (kDa). Note: The specks seen in (B) was only seen in the green channel and therefore did not interfere with the quantification of FAAH or GAPDH.

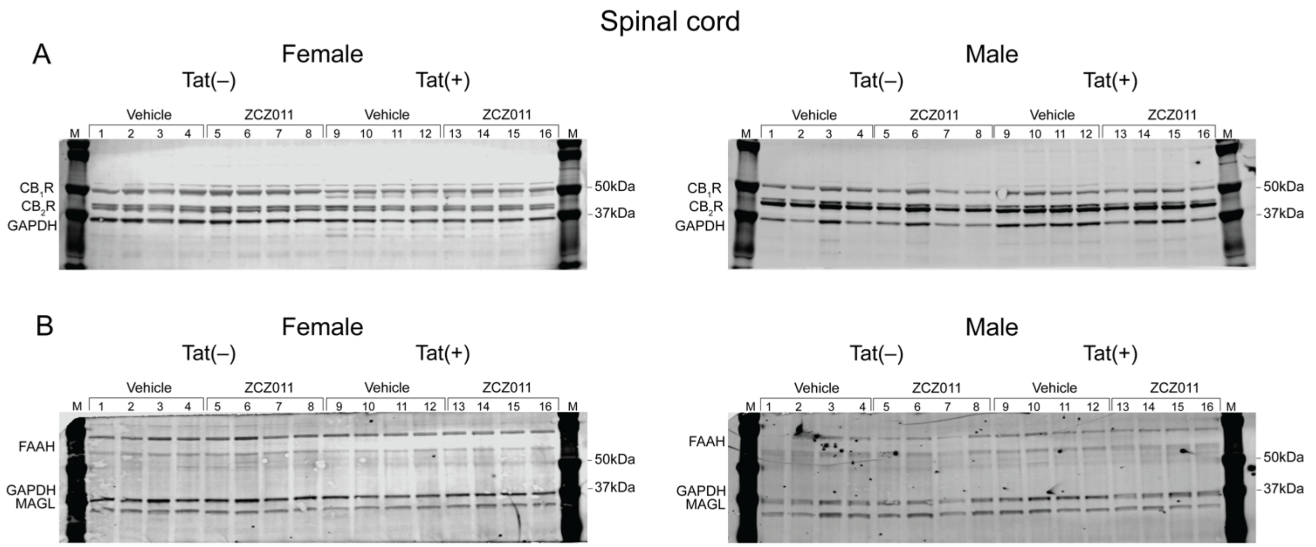

**S7_Fig. Original and unedited blots of CB$_1$R and CB$_2$R expression levels in the spinal cord.** Images show original (**A**) CB$_1$R, CB$_2$R, and GAPDH and (**B**) FAAH, MAGL, and GAPDH for females and males. Tat(–) vehicle- and ZCZ011-treated mice are represented by lanes 1-4 and 5-8 respectively. Tat(+) vehicle- and ZCZ011-treated mice are represented by lanes 9-12 and 13-16 respectively. M: molecular weights of marker protein (kDa).
